# Understanding the biochemical significance of the mitochondrial pyruvate carrier in the rodent malarial parasite, *Plasmodium berghei*

**DOI:** 10.1101/2025.01.06.631440

**Authors:** Nivedita Pandey, Chitralekha Sen Roy, Asutosh Bellur, Hemalatha Balaram

## Abstract

Mitochondrial pyruvate carrier (MPC) protein has been established as the key pyruvate transporter across the inner mitochondrial membrane. In *Plasmodium*, flux through the tricarboxylic acid (TCA) cycle is low in the asexual stages that is upregulated in the gametocytes with greater utilization of glutamine than glucose to drive the cycle. Acetyl-CoA in the mitochondrion has been established to be essential for the parasite. In this context, we established the biochemical function of *P. berghei* MPC as a pyruvate transporter through functional complementation in yeast. In *P. berghei*, both subunits, MPC1 and MPC2 were found to localize to the mitochondrion and surprisingly, were found to be non-essential for the blood stages and for ookinete formation. Through metabolite tracing studies using ^13^C_6_-glucose and ^13^C_5_^15^N_2_ L-glutamine, we have derived the biochemical basis for this observed phenotype of Δ*mpc1Δmpc2* parasites. Our studies suggest the presence of an aminotransferase localized to the mitochondrion that converts alanine to pyruvate, leading to the continued production of acetyl-CoA in the organelle even in the absence of MPC. This highlights the existence of metabolic plasticity, a survival strategy, that enables the utilization of different metabolic precursors for the generation of acetyl-CoA in the mitochondrion.

## Introduction

Pyruvate, being present at the branch point of glycolysis, gluconeogenesis, and TCA cycle, regulates carbohydrate, fatty acid, and amino acid metabolism (Zangari et al., 2020). Pyruvate oxidation through the TCA cycle results in the production of the reductants NADH and FADH_2_, which are then utilised during oxidative phosphorylation for energy generation (Martínez-Reyes and Chandel, 2020; Voet et al., 2016). Although biochemical studies published in 1971 (Papa et al., 1971) on the uptake of ^14^C pyruvate by rat liver mitochondria suggested the involvement of a translocator, it was only in the year 2012 that the genes for this transporter, Mitochondrial Pyruvate Carrier (MPC) were identified and characterized from *S. cerevisiae*, *Drosophila,* and human (Bricker et al., 2012; Herzig et al., 2012). Through their studies, it is now well known that MPC is located on the inner mitochondrial membrane and is composed of two subunits of small molecular mass proteins, MPC1 and MPC2. These two subunits assemble to form a stable and functional heterodimer for the import of pyruvate molecules into the mitochondrial matrix whereas though, individual subunits can also form homodimers, they are functionally inactive (Tavoulari et al., 2022, 2019). Studies have shown that any impairment in MPC function causes dysregulation of pyruvate metabolism leading to numerous metabolic diseases such as mitochondrial disorders, cancer, diabetes, and neurodegenerative disorders (Bender and Martinou, 2016; Chai et al., 2019; Disease et al., 2016; Divakaruni et al., 2017; Li et al., 2016). The function of MPC in regulating pyruvate transport activity has been examined in different organisms (Lyu et al., 2023; Negreiros et al., 2021; Štáfková et al., 2016).

Acetyl-CoA, a central metabolite also represents a crucial branch point in cellular metabolism due to its involvement in diverse metabolic pathways such as the TCA cycle, amino acid metabolism, fatty acid biosynthesis, and protein acetylation (Pietrocola et al., 2015; Shi and Tu, 2015). From studies reported thus far, there are three routes through which acetyl-CoA molecules are produced in *Plasmodium*. Apart from the mitochondrion and the apicoplast, organelles that generate acetyl-CoA pools (Chan et al., 2013; Foth et al., 2005; Nair et al., 2023; Oppenheim et al., 2014; Seeber et al., 2008), there is another important route for synthesising acetyl-CoA (Prata et al., 2021). The enzyme acetyl-CoA synthetase (ACS) located within the cytosol and the nucleus, catalyses the conversion of acetate into acetyl-CoA in an ATP-dependent manner (Summers et al., 2022; Prata et al., 2021). Recent studies on the role of acetyl-CoA pools in the regulation of diverse cellular processes in *Plasmodium* have drawn significant attention towards the discovery of new effective antimalarial (Jeffers and Child, 2022; Prata et al., 2021; Summers et al., 2022). Studies by Prigge and co-workers have established an important role for the mitochondrion in *Plasmodium*, as a major source of acetyl-CoA for the acetylation of the proteins present in both the nucleus and cytoplasm, thus suggesting the significance of mitochondrial acetyl-CoA in controlling a broad range of cellular functions and metabolic pathways (Nair et al., 2023). Mitochondrial branched-chain ketoacid dehydrogenase (BCKDH) and alpha-ketoglutarate dehydrogenase (KDH) complexes in *Plasmodium* have been shown to catalyse the conversion of pyruvate into acetyl-CoA (Chan et al., 2013; Oppenheim et al., 2014). MPC which mediates the entry of glucose-derived pyruvate into the mitochondrial matrix, may also play an essential role within the malarial parasite in the maintenance of the pool of mitochondrial acetyl-CoA and TCA cycle flux.

A recent deposition in bioRxiv describes studies on *P. falciparum* MPC (Rajaram et al., 2024). In this paper, we report our studies on the MPC of the rodent malaria parasite, *P. berghei*. To begin with we have established the biochemical function of *P. berghei* MPC as a pyruvate transporter through a functional complementation assay in yeast. This was followed by confirmation of the localization of both MPC1 and MPC2 subunits to the parasite mitochondrion. The essentiality of *Pb*MPC1 and *Pb*MPC2 subunits was examined using gene knockout strategies and the impact of MPC deletion on the growth phenotype and fitness was studied. Our results show that MPC1 and MPC2 subunits are dispensable for asexual blood stage growth, and the successful *in vitro* formation of ookinetes indicates that *in vitro* activation of gametocytes into gametes followed by fertilization and zygote formation is also unimpaired. Isotope tracing using ^13^C_6_-glucose and ^13^C ^15^N L-glutamine, and LC-MS, to address the complete absence of any phenotypic defects indicated the existence of metabolic plasticity in the murine malaria parasite that possibly enables mitochondrial acetyl-CoA production from alanine through an aminotransferase that interconverts pyruvate and alanine.

## Experimental procedures

### *Phylogenetic* analysis of *P. berghei* MPC proteins

*P. berghei* genes PBANKA_1354200 and PBANKA_1333600 annotated as putative MPC1 and MPC2, respectively were retrieved from the *Plasmodium spp.* functional genomics database, PlasmoDB (http://PlasmoDB.org) (Aurrecoechea et al., 2009). A total of 122 orthologs of MPC1 and MPC2 protein sequences from human, infectious fungi, *Drosophila melanogaster*, *Arabidopsis thaliana*, Kinetoplastids, Apicomplexans and Dinoflagellates were retrieved upon submitting *Saccharomyces cerevisiae* MPC1 or MPC2 sequences as queries into BLASTp in EuPathDB (http://eupathdb.org) and NCBI BLAST database. The multiple sequence alignments were performed using a built-in MUSCLE program in MEGA11 software (Tamura et al., 2021) and an outgroup-rooted phylogenetic tree was constructed using the maximum likelihood method and Le Gascuel’s model (Quang et al., 2008). The reliability of the phylogenetic tree was evaluated by the bootstrap method (N=500 replicates) and bootstrap values less than 50 were filtered out. Initial tree(s) for the heuristic search were obtained automatically by applying Neighbor-Join and BioNJ (Gascuel., 1997) to a matrix of pairwise distances estimated using the JTT model (Jones et al., 1992) and then selecting the topology with superior log likelihood value. This analysis involved 122 amino acid sequences. There was a total of 467 positions in the final dataset.

### *P. berghei* culturing and experimental animals

*P.berghei* ANKA strain was used in the study. 6-8 weeks old of either male or female BALB/c or C57BL/6 mice were routinely used as host for the maintenance of *P. berghei* parasites. The percent parasitemia was estimated by microscopic examination of a Giemsa-stained smear of blood collected from a tail snip. For transfection of parasites with DNA, standard protocols were used (Nagappa et al., 2019). Male or female C57BL/6 mice were used as host for the generation of single Δ*mpc1, Δmpc2,* and double Δ*mpc1Δmpc2* knockout lines and for carrying out growth rate experiments across intra-erythrocytic asexual stages and, for enumeration and collection of gametocytes (sexual). All animal experiments with BALB/c and C57BL/6 mice were conducted following standard operating procedures approved by the Institutional Animal Ethics Committee (IAEC) of JNCASR for Prevention of Cruelty to Animals Act of 1960 and Breeding and Experimentation Rules of 1998, Constitution of India.

### Generation of *P.berghei* MPC gene knockout and tagging constructs for transfection

The library clones procured from PlasmoGEM repository (Wellcome Trust Sanger Institute, UK) (Gomes et al., 2015; Schwach et al., 2015) for *P. berghei* MPC1 [library clone id: 2435h06] and MPC2 [library clone id: c19eo7] were first validated by PCR and restriction digestion. The procedures followed for the generation of knockout and tagging constructs were as described earlier (Gomes et al., 2015; Pfander et al., 2011). The generated MPC1 and MPC2 final knockout constructs were subjected to NotI digestion and purified prior to transfection. For tagging of the endogenous copy of *mpc1* and *mpc2* genes in *P. berghei* with GFP, the C-terminal GFP-DDD-1XHA (DDD: DHFR degradation domain) tagging constructs of MPC1 and MPC2 were generated using the *in vivo* recombineering method as previously described (Gomes et al., 2015). Briefly, the bacterial selection marker (Zeo-PheS) was first inserted immediately upstream of the stop codon, without modifying the open reading frame of the *mpc1* or *mpc2* gene that is to be tagged. The Zeo-PheS cassette was replaced by the GFP-DDD-1X HA tag, introduced in frame at the C-terminus using Gateway LR Clonase (Invitrogen). The sequences of the oligonucleotides used for the generation of knockout and tagging constructs and PCR validation are provided in Supplementary Table 1.

### Transfection of *P. berghei*, genotyping of drug-resistant parasites and cloning by limiting dilution

Established protocols were followed for transfection of *P. berghei* (Janse et al., 2006). Briefly, 5-8 μg of DNA was mixed with schizont-enriched parasites and 100 μl of P5 nucleofector solution (Lonza). Transfection was done using Amaxa 4D-nucleofector system (Lonza), using the program ‘FP167’ or ‘FP148’, followed by injection of transfected parasites into two mice. Drug selection was started 48 hrs post-infection by feeding the mice *ad libitum* with pyrimethamine (70 μg ml^-1^) dissolved in water. The drug-resistant parasites were enumerated by microscopic examination of Giemsa-stained smears of tail blood. At ∼10 % parasitaemia, parasites were harvested, glycerol stocks made and genomic DNA isolated. The confirmation of Δ*mpc1, Δmpc2,* and Δ*mpc1Δmpc2* parasites and, MPC1-GFP-DDD-1XHA and MPC2-GFP-DDD-1XHA tagged parasites was achieved by PCR using appropriate oligonucleotide primers. The sequences of oligonucleotides used for PCR genotyping are provided in supplementary Table S1. Thereafter, *in vivo* cloning by limiting dilution was performed following standard procedures (Orr et al., 2012) to obtain clonal lines for single Δ*mpc1, Δmpc2, and* double Δ*mpc1Δmpc2* knockouts before phenotypic analysis across different stages of the *P. berghei* life cycle was carried out.

### Generation of marker-free ***Δmpc2*** parasites

A widely used negative selection protocol was adopted on *Δmpc2* parasites to remove the human dihydrofolate reductase and a yeast bifunctional protein that combines yeast cytosine deaminase and uridyl phosphoribosyltransferase (hDHFR-yFCU) cassette by using 5-fluorocytosine (5FC) as the selection drug (Manzoni et al., 2014). A glycerol stock of *Δmpc2* parasites was injected into 2 C57BL/6 mice. At parasitaemia between 0.5-1 %, 5FC (1.5 mg ml^-1^) in drinking water was administered *ad libitum* to mice. A noticeable drop in parasitaemia was observed after 48 hrs and the re-appearance of the parasites under 5FC drug pressure was monitored regularly through the microscopic observation of Giemsa-stained smears. At ∼10 % parasitaemia, parasites were harvested, genomic DNA isolated and the absence of the drug-selection cassette, hDHFR-yFCU was confirmed by PCR. The marker-free *Δmpc2 P. berghei* clonal lines were obtained by the *in vivo* limiting dilution cloning (Orr et al., 2012). In addition, loss of resistance to pyrimethamine due to excision of the hDHFR marker was confirmed by injecting marker-excised *Δmpc2 P. berghei* line into BALB/c mice and upon parasitaemia reaching ∼1-1.5 %, drug pressure (pyrimethamine 70 μg ml^-1^) was applied and continued for 25 days. The drug-sensitive *Δmpc2* parasite line was used further for genetic manipulation for the second time to generate the Δ*mpc1Δmpc2* parasite line.

### Asexual growth rate, enumeration of gametocytes, and *in vitro* ookinete conversion

To assess the growth rate during asexual blood stages, glycerol stocks of wildtype, Δ*mpc1, Δmpc2,* and Δ*mpc1Δmpc2* parasites were injected independently into C57BL/6 mice and parasitaemia monitored by microscopic examination of Giemsa-stained blood smears. The blood was harvested at parasitaemia between 0.5-1% and 10^4^ parasites of wild type and knockout lines were injected into C57BL/6 mice, five in each group (N=5). Post-infection, parasitaemia was monitored daily for each genotype by microscopic counting of parasites in Giemsa-stained blood smears. The percent survival curve among the 5 mice infected with either wildtype or the Δ*mpc* knockout lines was plotted with the X-axis representing time (number of days) and the Y-axis the % survival rate.

To quantify the production of gametocytes, mice were first injected intraperitoneally with phenylhydrazine (12.5 mg ml^-1^ dissolved in 0.9 % NaCl). 4 mice were used for each line of *P. berghei*. Two days thereafter, 10^5^ parasites of wild type, Δ*mpc1, Δmpc2,* and Δ*mpc1Δmpc2* were injected into the mice. Two days post-infection, all mice were fed *ad libitum* with sulfadiazine (30 mg L^-1^ in drinking water) containing drinking water. On day 3 post-sulfadiazine treatment, Giemsa-stained smears of tail vein blood were made for each genotype, and gametocytes were enumerated. The female-to-male gametocyte ratio was also calculated and plotted.

For *in vitro* ookinete conversation assay, C57BL/6 mice were intraperitoneally injected with phenylhydrazine (12.5 mg ml^-1^). After 2 days, 10^7^ parasites of each wild type, Δ*mpc1,* Δ*mpc2* and Δ*mpc1Δmpc2* were injected independently into mice. On day 3 post-infection, parasitaemia was monitored by microscopic examination of Giemsa-stained blood smears, and sulfadiazine (30 mg L^-1^ in drinking water) was fed ad libitum to the animals. Two days after sulfadiazine treatment, mice were bled and 350 µl infected blood containing gametocytes was incubated in 10 ml ookinete medium (RPMI 1640, 25 mM HEPES, 370 µM hypoxanthine, 20 % fetal bovine serum, 0.2 % sodium bicarbonate, 100 μM xanthurenic acid and 40 mg L^-1^ gentamycin, pH 8.3) at 1:25 dilution for 24 hours at 19 ℃. Thereafter, the cell suspension was centrifuged, Giemsa-stained blood smears were examined and percent ookinete conversion was quantified using the formula [(number of ookinetes observed)/ (total number of female gametocytes, zygotes and ookinetes) × 100 % (Patil et al., 2020).

### Live-cell microscopy

Blood from mice infected with *P. berghei* lines expressing either GFP-tagged *Pb*MPC1 or *Pb*MPC2 was harvested in incomplete RPMI medium containing heparin (1mg ml^-1^). The cells were then washed three times with 1X PBS and recovered by centrifugation at 900 xg for 5 minutes. For co-localization studies, mitochondrial staining was performed. Briefly, the cells were resuspended in incomplete RPMI 1640 medium containing 120 nM MitoTracker^TM^ Orange CMTMRos (Invitrogen^TM^) and incubated at 37 ℃ in a candle jar for 25 minutes and then Hoechst 33342 (10 µg ml^-1^) was added and incubated for additional 10 minutes at 37 ℃ in a candle jar. Thereafter, the cells were washed three times with pre-warmed 1x PBS and recovered by centrifugation at 900 xg for 5 minutes. Finally, the cells were resuspended in VECTASHIELD mounting medium (Vector laboratories) and mounted on poly-L lysine coated glass slides using a coverslip and sealed. The images were acquired within 40 minutes using Delta Vision microscope (GE Healthcare) under oil immersion objective at 100x magnification. For evaluation of co-localization, image processing was performed, and Pearson correlation coefficient (PCC) values were calculated using Fiji software (Schneider et al., 2012).

### RNA isolation, cDNA synthesis and reverse transcription (RT)-PCR

Blood was drawn from infected mice at ∼10 % parasitaemia in incomplete RPMI 1640 medium containing heparin (1mg ml^-1^). The parasites were released from the erythrocytes by resuspension in erythrocyte lysis buffer (155 mM NH_4_Cl, 12 mM NaHCO_3_ and 0.1 mM EDTA) and incubation for 5 minutes at 4 ℃. The parasite pellet was harvested by centrifugation at 900 xg for 10 minutes. RNA was isolated using Trizol reagent and standard procedures (Nagappa et al., 2019). The RNA pellet was dissolved in 10-20 µL of DEPC treated water and stored at −80 ℃. Prior to cDNA synthesis, the RNA was treated with DNase I (NEB).

For first-strand cDNA synthesis, 1 µg of DNase I treated RNA was mixed with 625 µM dNTPs and 2.5 µM gene-specific reverse primer and the volume was made up with 0.1% diethylpyrocarbonate (DEPC) treated water. This was heat denatured for 5 minutes at 65 ℃ and placed on ice. Thereafter, RNase inhibitor (human placenta, NEB) and M-MuLV reverse transcriptase (NEB) were added and incubated for 1 hour at 45 ℃. Thereafter, the reaction mix was first incubated at 65 ^0^C for 20 minutes to heat inactivate M-MuLV reverse transcriptase and the newly synthesized cDNA was used as template in PCRs.

### Cloning of *mpc* genes into yeast expression plasmids

All constructs were generated in DH5α strain of *Escherichia coli* cells. The amplified fragments were cloned into yeast shuttle expression vectors, p413-GPD and p416-TEF for constitutive gene expression. The cDNA, synthesized from total RNA isolated from *P. berghei* was used as template in PCRs for amplifying the Pb*mpc1* and Pb*mpc2* genes. The PCR fragment of Pb*mpc1* gene with 6xHis tag at the 3′ end was cloned into p413-GPD vector using the restriction sites, EcoRI and SalI. *Pb*MPC2 gene with a 6xHis tag at the 3′ end was cloned into p416-TEF plasmid at BamHI and SalI sites. Likewise, *S. cerevisiae mpc1* and *mpc2* genes were amplified by PCR using yeast genomic DNA as template and cloned with a C-terminal 6xHis tag into p413-GPD (Sc*mpc1*) and p416-TEF (Sc*mpc2*). All clones were confirmed by DNA sequencing. The details of the oligonucleotide sequences used in the generation of the constructs are provided in Table S1.

### Transformation of yeast strains and spot assays

*Δmpc1, Δmpc2Δmpc3 and Δmpc1Δmpc2Δmpc3* yeast strains of genotype W303a MATa his3 lys2 met15 trp1 ura3 were obtained as a generous gift from Prof. Jarred Rutter, University of Utah, USA and characterized by PCR and spot growth assays (Supplementary Fig. S1). Yeast cells were transformed using the lithium acetate method (Daniel Gietz and Woods, 2002). Briefly, a single colony was inoculated into 5 mL of YPD growth medium and grown for 12 hours at 30 ^0^C at 200 rpm. The following day, the OD_600_ of the overnight culture was measured and inoculated into 5ml of YPD medium to yield OD_600_ of 0.2. This was further incubated for 3-5 hours to produce a culture with an OD_600_ of ∼0.4-0.6. Thereafter, the cells were pelleted down, washed two times with sterile MilliQ water and resuspended in a transformation mix consisting of 50% PEG, 1 mg mL^-1^ denatured salmon sperm DNA, 1M lithium acetate and 1.5-3 µg of plasmid DNA. The mixture was vortexed briefly and then kept in a 42 ^0^C water bath for 40 min for heat shock. Thereafter, cells were pelleted down at 900 xg for 5 minutes, washed two times with sterile MilliQ water and then plated on a solidified selective medium. mpc1Δ, mpc2Δmpc3Δ and mpc1Δmpc2Δmpc3Δ strains were co-transformed with two plasmids p413-GPD (His3 selection marker) and p416-TEF (Ura3 selection marker), either empty or having individual MPC genes. The co-transformed mpc1Δ, mpc2Δmpc3Δ and mpc1Δmpc2Δmpc3Δ yeast cells were grown in SD growth medium containing all amino acids but lacking histidine and uracil (Ura) for ensuring plasmid maintenance.

For spot assays, single colonies were picked up from the plate and inoculated into 5 mL of SD without His-Ura with the branched-chain amino acids leucine (150 mg L^-1^) and valine (100 mg L^-1^) in the medium. The culture was grown for 16-18 hours at 30 ^0^C at 200 rpm. The cells were pelleted down at 900 xg for 5 minutes and then washed twice with sterile MilliQ water. OD_600_ was checked and then the cells were diluted to an OD_600_ = 1 and serial 10-fold dilutions were spotted on agar plates containing SD-His-Ura medium with or without leucine and valine. Plates were incubated for 48 hours at 30 ^0^C before photographic capture of the image of the plates.

### Extraction of labelled metabolites

To obtain gametocyte-enriched *P. berghei* parasites for metabolite isolation, a glycerol stock of either wild-type or Δ*mpc1*Δ*mpc2* parasites was injected into a C57BL/6 mouse. The appearance of the parasites was monitored using Giemsa-stained smears of blood collected from tail snips. At parasitaemia between 1-2 %, 500 μL of infected blood was collected in 350 μL of RPMI medium containing heparin (1mg ml^-1^). 200 μL of this harvested blood containing *P. berghei* parasites was then passaged to 3-5 C57BL/6 mice pre-treated for two days with phenyl hydrazine to enhance the population of reticulocytes in the blood. 3-4 days post-infection, at a parasitemia of 10-15 %, mice were fed *ad libitum* for the following 2-3 days with sulfadiazine (3 mg in 100 mL drinking water) to kill all asexual blood stages. Thereafter, the infected blood having only gametocyte-stage parasites was harvested from mice in a tube with pre-warmed RPMI medium containing heparin (1mg ml^-1^), and the tube was immediately kept at 37 ℃.

The remaining procedures were performed in an incubator at 37 ℃ and the required materials (like tips, tubes, T-25 flasks, and media) were also pre-warmed to 37 ℃. The maintenance of the temperature at 37 ℃ was strictly ensured to prevent the activation of gametocytes into gametes. The gametocyte medium was used for the steps of leukocyte removal from whole blood and for washing the gametocytes prior to their *in vitro* culturing. Leukocytes were removed from whole blood using a cellulose column following published procedures (Venkatesan et al., 2012). The blood collected from the cellulose column was microscopically examined to confirm the absence of leukocytes. The next step involved separation of the gametocyte-enriched parasites from uninfected erythrocytes by density gradient centrifugation on 55 % Nycodenz in 1X phosphate buffered saline (PBS) using standard procedures (Rodríguez et al., 2002). The harvested fraction enriched in gametocytes was washed thrice with 5 mL of gametocyte medium and centrifuged at 2000xg for 5 minutes at 37 ℃. The enrichment of gametocytes was verified by microscopic examination of Giemsa-stained smears of the resuspended cells. For gametocyte counting, the gametocyte pellet was resuspended in 1 mL of gametocyte medium, and appropriate dilutions were made. The number of gametocytes per μL of the diluted blood was estimated using a Neubauer hemocytometer. The volume corresponding to 2 × 10^7^ gametocytes was added to a T25 flask containing 5 mL of minimal medium, incubated with gentle shaking at 37 ℃, 60 rpm, and parasites were allowed to pre-equilibrate *in vitro* for 90 minutes. The compositions of the gametocyte medium and minimal medium are provided in the supplementary section.

Metabolite labelling studies with wild-type or *Δmpc1Δmpc2 P. berghei* gametocytes were performed using ^13^C_6_-glucose and ^13^C_515_N_2_ L-glutamine. The pre-equilibrated gametocytes were washed with 5 mL of minimal medium without L-glutamine or glucose and thereafter incubated with either 2 mM ^13^C ^15^N L-glutamine or 8 mM U^13^C-glucose in 5 mL of minimal medium for 1 hr 30 minutes at 37 ℃, with gentle shaking at 60 rpm.

Control experiments were conducted with uninfected erythrocytes. 2 × 10^7^ erythrocytes were cultured for 90 minutes in the presence of either U-^13^C_6_-glucose or U-^13^C_5_^15^N_2_ L-glutamine under conditions as described for the parasites. For wild-type and *Δmpc1Δmpc2* parasites, a total of two independent biological replicates (N=2) of the experiment were carried out. In each experiment of either wild-type or *Δmpc1Δmpc2* gametocytes, 2 technical samples were prepared in two T-25 flasks under the conditions mentioned above.

Post incubation with the isotope enriched glucose or glutamine, the cells were harvested by centrifugation at 2000 xg for 5 minutes. This step was carried out at 37 ℃ to avoid gametocyte activation. The cells were resuspended in 1 mL of ice-cold 1X PBS, pH 7.4 (prepared in LC-MS grade water) and centrifuged at 10,000 rpm for 2 minutes at 4 ℃ to eliminate contamination from the spent medium. To extract the metabolites, 1 mL of ice-cold 90 % methanol (LC-MS grade) was added. Parasites were rapidly resuspended by vortexing and 10 μL of a mixture of internal standards containing 30 μM PIPES (1,4-Piperazinediethanesulfonic acid) and 15 μM CAPS (N-cyclohexyl-3-aminopropanesulfonic acid) was added. The extract was centrifuged at 13,000 rpm for 10 minutes at 4 ℃. The supernatant was transferred to a fresh, pre-chilled Eppendorf tube, dried immediately under nitrogen flow, and stored at −80 ℃.

### LC-MS Data acquisition

Metabolites were analysed by LC-MS using hydrophilic interaction liquid chromatography (HILIC) and Orbitrap-MS. The dried extracts of metabolites were dissolved in 30 μL of 40 % acetonitrile (LC-MS grade) and vortexed for 20 minutes at 4 ℃. The sample was then centrifuged at 13,000 rpm for 15 minutes at 10 ℃. 4 μL of the sample was injected into a ZIC-pHILIC column (5 μm, 150 × 2.1 mm; SeQuant, Merck) attached to Dionex Ultimate 3000 chromatography system coupled to Orbitrap Q Exactive HF (Thermo Scientific). A guard column (20 × 2.1 mm; SeQuant, Merck) was connected before the ZIC-pHILIC column. 20 mM ammonium carbonate, pH 9.4 (A), and 100% acetonitrile (B) were used as the mobile phases. A gradient run, adopted from a study by Giannangelo and co-workers (Giannangelo et al., 2020) was modified further for our experiments. This involved a 45-minute gradient starting from 80% B for the first 1 minute, then 80% to 40% B in 20 minutes, 40% to 5% B in 10 minutes and held constant at 5% B for 3 minutes for washing, and re-equilibration of the column at 80% B for 10 minutes. For the MS analysis, Orbitrap Q Exactive HF (Thermo Scientific) equipped with a heated electrospray ionization source was operated in dual polarity mode (positive and negative) using the settings described previously (Creek et al., 2011; MacKay et al., 2015) with minor modifications; mass resolution of 60000 from m/z 70-1050, sheath gas flow rate 30, aux gas flow rate 15, spray voltage 4 kV in positive polarity and 3.5 kV for negative polarity, capillary temperature 320 ℃, auxiliary gas heater temperature 100 ℃, AGC target 1 × 10^6^, maximum IT 200 ms, micro scans 3. The flow rate was set at 0.1 ml/minute, the column temperature at 45 ℃, and the autosampler was maintained at 10 ℃. The instrument was calibrated at regular intervals and conditioning of the column was done before each batch of samples by running blanks and a mixture of reliable standards (26 metabolites). Metabolites and their isotopically labelled counterparts were identified manually based on its retention time and extracted ion chromatograms (EICs) using Qual Browser (Xcalibur 4.1, Thermo Scientific). Quality control (QC) samples were also analysed periodically and samples within each batch were randomized to avoid any systemic drift on metabolite signals.

### Data processing and analysis of LC-MS metabolomics experiments

All metabolomics data were analysed using EI-MAVEN, 0.12.1 version software (Agrawal et al., 2019). Briefly, it involved the conversion of a raw data file to mzXML format using the msConvert tool (Chambers et al., 2012). The converted raw file was then loaded onto EI-MAVEN software. Metabolites were identified by using a pre-entered selected list of target compounds with m/z values and retention times which enabled EI-MAVEN to obtain extracted ion chromatograms (EIC) of various metabolites (extracted ion chromatogram) from the MS1-full scan data of each sample. The peak areas of the identified metabolites were recorded and exported as an Excel sheet in CSV format as described by Agrawal et al., 2019.

For all ^13^C_6_-glucose or ^13^C ^15^N L-glutamine labelled samples, the peak area of each isotopologue of a metabolite was first normalized to the 2 × 10^7^ gametocytes and the correction for the natural abundance (NA) of isotopes within a metabolite was performed using an R-based tool, IsoCorrectoR (Heinrich et al., 2018). Post-NA correction, the percentage labelling of each isotopologue of a metabolite was calculated from each sample of either wild-type or *Δmpc1Δmpc2* parasites and then plotted on the graph. The samples prepared from two independent biological replicates carried at least two technical replicates each. The error bars on the graph indicate the standard deviation (SD). In ^13^C_5_^15^N_2_ L-glutamine labelled *Δmpc1Δmpc2* samples, only one sample from each biological replicate could be prepared. Significance was determined by using unpaired t-test with Welch’s correction, with significant differences, p-value < 0.05, p-value < 0.01, p-value < 0.001, p-value < 0.0001 were indicated by one, two, three, and four asterisks, respectively.

## Results

### MPC in Plasmodium

The genes annotated as putative MPC1[Gene ID: PBANKA_1354200] and MPC2 [Gene ID: PBANKA_1333600] subunits in *P. berghei* were retrieved from PlasmoDB (Aurrecoechea et al., 2009). Both *mpc1* and *mpc2* are located on chromosome 13 with only one intron in the *mpc1* gene and three introns in *mpc2* gene sequence. The transcription of *mpc1* and *mpc2* genes in *P. berghei,* yielding transcripts of the expected size was confirmed by reverse transcription (RT)-PCR (Figure 1A).

**Figure 1.**
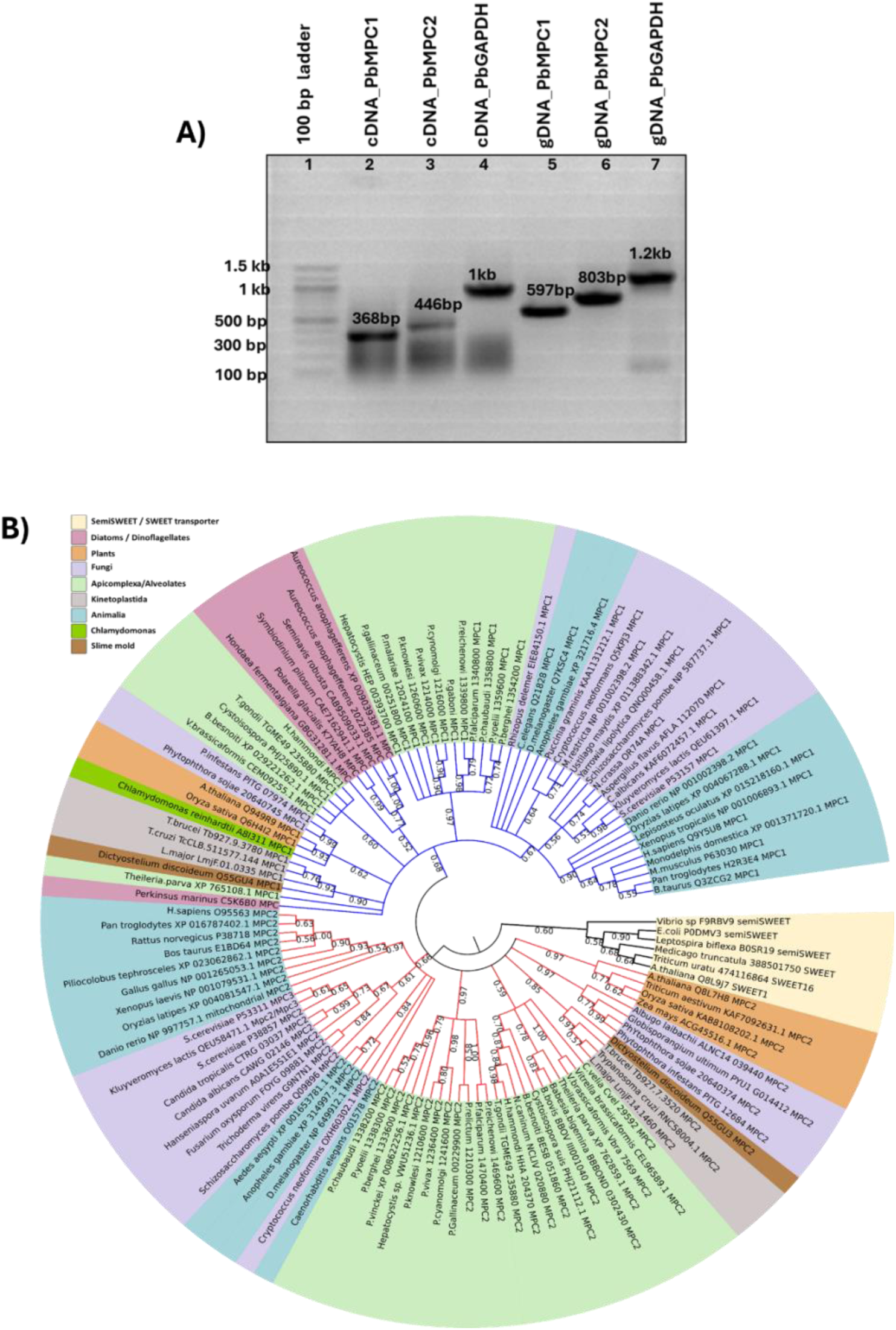
MPC1 and MPC2: expression in *P. berghei* and phylogenetic relationship. A) Confirmation of *mpc1* and *mpc2* gene expression in *P. berghei* by RT-PCR. Lane 1, 100 bp DNA size marker; lanes 2 – 4, RT-PCR with RNA from *P.berghei* ANKA parasites, lanes 5 – 7, PCR with parasite genomic DNA (gDNA) as template using gene-specific Pb*mpc1*, Pb*mpc2* and Pb*gapdh* forward and reverse primers. Pb*gapdh* gene was used as a positive control for RT-PCR as well for PCR with genomic DNA as template. kb is kbp (kilo base pairs). B) Phylogenetic tree of MPC sequences from diverse organisms. The maximum likelihood method and Le Gascuel’s (2008) model were chosen to represent and analyze the evolutionary history from 500 bootstrap replicates. Branches corresponding to partitions reproduced in less than 50% of bootstrap replicates are collapsed. Clades corresponding to major kingdoms/phyla are highlighted in different colours. The protein sequences of Semi-SWEET / SWEET transporters (highlighted in yellow shade) served as an “outgroup” for constructing a rooted-phylogenetic tree.

DeepMito (http://busca.biocomp.unibo.it/deepmito) predicts the localization of both the subunits *Pb*MPC1 (score: 0.8) and *Pb*MPC2 (score: 0.81) to the mitochondrial inner membrane. The protein sequences of MPC1 and MPC2 from various *Plasmodium* species share 68% to 93% sequence identity, respectively suggesting their conserved function. The multiple protein sequence alignments of MPC1 and MPC2 orthologs (from *P. berghei*, human, mouse, *Drosophila*, *S. cerevisiae)* highlight the presence of invariant and conserved residues indicating their possible roles in structure, subunit assembly and substrate binding (Supplementary Figure S2).

A phylogenetic tree with a total of 122 MPC1 and MPC2 protein sequences was constructed using the maximum likelihood method and Le Gascuel’s model (Quang et al., 2008) (Figure 1B). The phylogenetic tree was rooted by including the protein sequences of semiSWEET / SWEET transporters as an “outgroup”. These transporters have been suggested to be distant homologs of MPCs, with their protein sequences sharing a very low identity of ∼10% with MPC (Xu et al., 2014). The two subunits of MPC were identified in almost all organisms belonging to the domain Eukarya while MPC identified in alveolates such as *Perkinsus marinus,* Stramenopiles (eg. *Aureococcus anophagefferens),* and Dinoflagellates (eg. *Symbiodinium pilosum)* had only a single fused MPC protein containing all the signature sequence motifs present in MPC1 and MPC2 subunits. The phylogenetic tree shows that MPC is divided into two major groups: MPC1 and MPC2. The MPC1 and MPC2 of *P. berghei* cluster with MPC1 and MPC2, respectively of other *Plasmodium* species. In contrast, MPC1 or 2 of *T. gondii* form a mini cluster with their closely related apicomplexan parasites; *Cystoisospora*, *Besnoitia,* and *Hammondia.* It should be noted that the length of the *Tg*MPC1 protein sequence is longer than MPC1 of *Plasmodium sp.* by about 50-60 residues. The tree also shows the divergence of *Plasmodium* MPCs from *Trypanosoma sp.* of the phylum Kinetoplastida. In yeast, MPC encodes for an additional MPC subunit, MPC3. The MPC3 subunit of *S. cerevisiae* groups together with MPC2 sequences as it shares a high sequence identity (∼80 %) with the yeast MPC2 subunit. We also observed that the single-fused MPC protein sequences from *alveolates* and *dinoflagellates* are positioned in the MPC1 cluster, which suggests that the MPC from these organisms shares greater sequence similarity with MPC1 than with the MPC2 subunit.

### Probing the pyruvate transport function of *Pb*MPCs through functional complementation in yeast

The ability of *P. berghei mpc1* (Pb*mpc1)* and *P. berghei mpc2 (*Pb*mpc2)* genes to transport pyruvate was examined by functional complementation in yeast strains lacking *mpc1* (ScΔ*mpc1*), *mpc2* and *mpc3* (ScΔ*mpc2*Δ*mpc3*) and, *mpc1, mpc2* and *mpc3* (ScΔ*mpc1*Δ*mpc2*Δ*mpc3*) (Bricker et al., 2012). Pyruvate serves as the precursor molecule for the biosynthesis of the branched-chain amino acids, leucine and valine. Yeast cells that lack MPC genes cannot grow in minimal medium lacking leucine and valine and this requirement becomes the basis of the functional complementation assay (Figure 2A). The MPC1 gene (Pb*mpc1* or Sc*mpc1*) with a C-terminal 6x His tag was cloned into the yeast expression plasmid, p413-GPD whereas the *mpc2* gene (Pb*mpc2* or Sc*mpc2*) with a C-terminal 6x His tag was cloned into yeast expression plasmid p416-TEF. *mpc* mutant yeast cells were then co-transformed with p413-GPD and p416-TEF plasmids either empty or expressing the individual MPC subunits. Spot assays were done on plates containing synthetic medium with or without the branched-chain amino acids, leucine and valine, and histidine and uracil were omitted to ensure plasmid maintenance.

**Figure 2.**
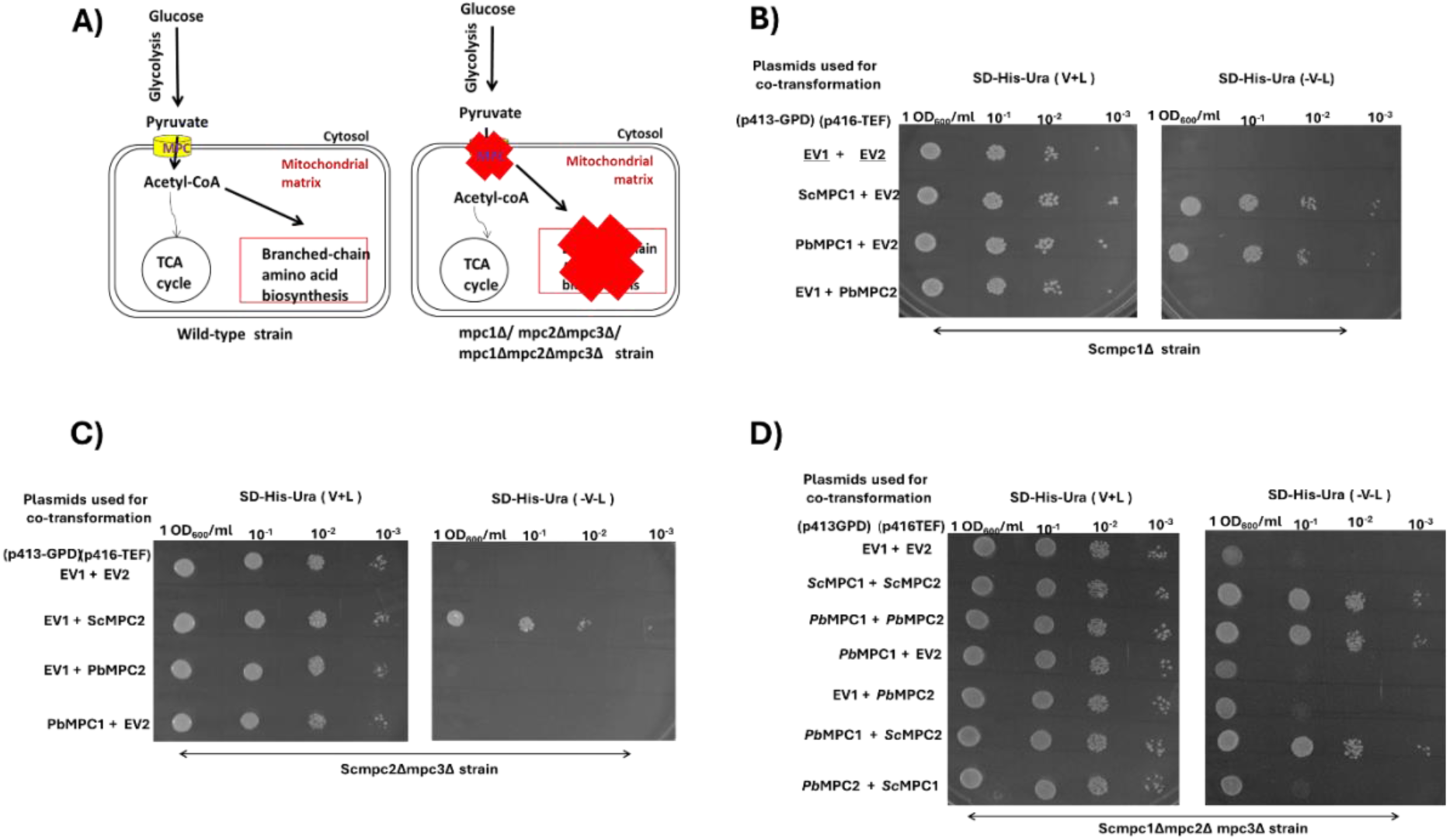
*P. berghei* MPC1 and MPC2 assemble into a functional complex that mediates the transport of pyruvate from the cytosol into the mitochondria. A) Schematic of the basis of the yeast functional complementation assay using *mpc* knockout strains of *S. cerevisiae*. Panels (B), (C), and (D) show results from spot assays with Sc*Δmpc1*, Sc*Δmpc2Δmpc3*, and Sc*Δmpc1Δmpc2Δmpc3* yeast strains, respectively. MPC-deficient yeast strains co-transformed with empty vectors, p413-GPD, and p416-TEF served as negative controls in all spotting assays. The co-transformed yeast cells were diluted to an OD_600_ = 1 and serial 10-fold dilutions were spotted on agar plates containing SD-His-Ura medium with (V+L, left subpanel) or without leucine and valine (-V-L, right subpanel). Plates were incubated for 48 hours at 30 ℃ before photographic acquisition of the images. Histidine and uracil were omitted from the synthetic dextrose (SD) medium to ensure plasmid maintenance. N=3 independent experiments were done.

To maintain uniformity in growth conditions, both p413-GPD and p416-TEF as empty vectors or carrying Sc or Pb *mpc* genes were co-transformed in all complementation assays. Transformation with empty plasmids served as negative controls. The growth defect of *Δmpc1* yeast cells in synthetic medium lacking Leu and Val could be rescued when transformed with either *Sc*MPC1 or *Pb*MPC1 expressing plasmids (Figure 2B). This indicates that the *Pb*MPC1 subunit complements the function of *Sc*MPC1 in *Δmpc1* yeast strain and the function of MPC1 subunit is conserved across the two species.

*Δmpc2Δmpc3* yeast strain was used to examine if *Pb*MPC2 can restore the growth defect and our results show that neither *Pb*MPC2 nor *Pb*MPC1 subunits complement the function of yeast MPC2 (Figure 2C). The inability of *Pb*MPC2 in rescuing the growth defect might be due to inability to form a functional complex with yeast MPC1 or structural differences in the complex of *Pb*MPC2-*Sc*MPC1 resulting in a non-functional heterocomplex, incapable of transporting pyruvate. The triple deletion *Δmpc1Δmpc2Δmpc3* yeast strain was used to examine if both the subunits of *Plasmodium* MPCs can rescue the growth defect in yeast cells lacking MPC genes and if the individual subunits of *Plasmodium* (*Pb*MPC1 or *Pb*MPC2) can form a functional homo-complex and mediate the uptake of pyruvate into the mitochondrial matrix. Yeast cells co-transformed with *Sc*MPC1 and *Sc*MPC2 together could rescue the growth defect whereas cells transformed with individual yeast subunits could not. In yeast, it has been reported earlier that both the subunits of MPC are necessary for mitochondrial pyruvate transport activity (Bricker et al., 2012; Herzig et al., 2012; Tavoulari et al., 2019). Our result indicates that individual subunits, either *Pb*MPC1 or *Pb*MPC2 are not sufficient but both subunits of *Plasmodium* MPC are required to form a functional heterocomplex. The yeast cells co-transformed with *Pb*MPC1 and yeast MPC2 expressing plasmids could also rescue the defect in the triple *mpc* knockout yeast strain indicating the conservation of structural features in the PbMPC1 subunit necessary for forming a functional heterocomplex. Unlike *Pb*MPC1, *Pb*MPC2 could not rescue the growth defect when co-transformed with yeast MPC1 but restored growth only when expressed together with *Pb*MPC1 (Figure 2D).

To confirm that the rescue seen is due to the expression of *Sc* and *Pb* MPCs, the expression of 6XHis tagged ScMPC1, ScMPC2, PbMPC1 and PbMPC2 genes in yeast was validated at the transcript level by RT-PCR and protein level by Western blotting. Our RT-PCR results confirmed the expression of Pb*mpc1* and Pb*mpc2* genes at the transcript level in *mpc1Δ* cells transformed with PbMPC1 or PbMPC2 expressing plasmids (Supplementary Figure. S3). Despite repeated attempts, we were unable to detect protein expression using anti-His antibodies of both *Sc* and *Pb* MPC1 and MPC2 subunits fused with a 6X-His tag at their C-terminus.

### Expression and sub-cellular localization of PbMPC1 and PbMPC2 in *P. berghei* parasites

To determine the sub-cellular localization of MPC1 and MPC2 subunits in *P. berghei*, we generated two independent transgenic parasite lines expressing either endogenous MPC1 or MPC2 fused to a GFP_DDD_1xHA (DDD: DHFR Degradation Domain) tag at their C-terminal ends. The strategy followed and validation of the constructs are shown in Supplementary Figure S4. PCR genotyping was carried out to confirm the successful generation of MPC1-GFP_DDD_1xHA and MPC2-GFP_DDD_1xHA tagged parasite lines. 5’ and 3’ integration PCRs resulted in expected fragments of 3.4 kb and 8.5 kb, respectively for MPC1, and 8.3 kb and 6kb, respectively for MPC2, thereby confirming the integration of the tag and selection cassette at the expected loci (Supplementary Figure S4). Live cell microscopy of parasites carrying GFP-tagged endogenous copy of MPC1 and MPC2 confirmed expression across all blood stages. Further, the GFP signal was found to overlap with that of MitoTracker orange CMTMRos dye, indicating their localization to the mitochondrion of the parasites (Figure 3). In a recent study, MPC2 of *P. falciparum* was found to be localized to the mitochondrion (Rajaram et al., 2024).

**Figure 3.**
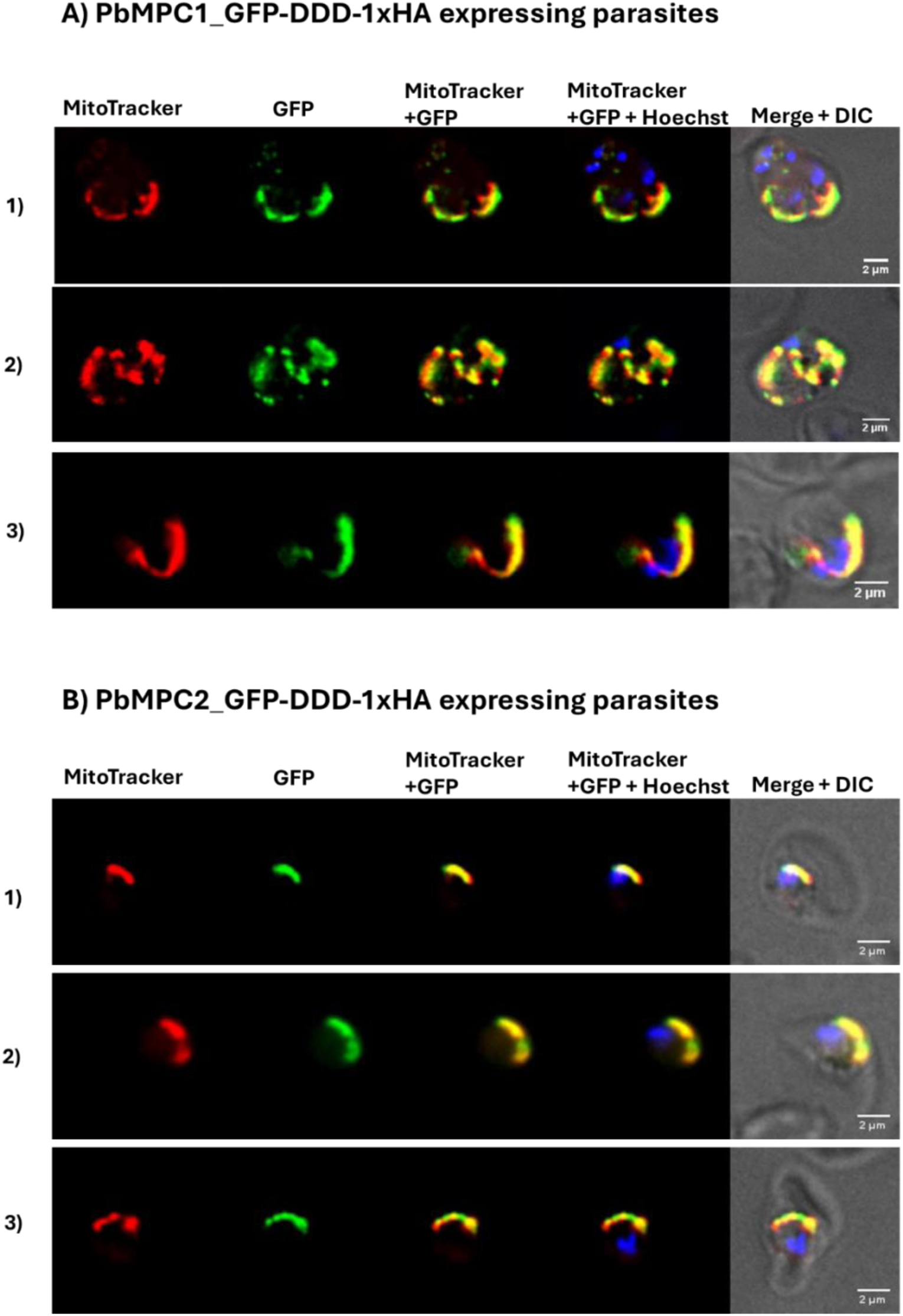
Expression and sub-cellular localization of MPC1 and MPC2 in the erythrocytic stages of *P. berghei.* A) and B) Live-cell microscopy images showing expression of PbMPC1-GFP-DDD-1XHA and PbMPC1-GFP-DDD-1XHA proteins in different intra-erythrocytic stages. In addition, the co-localization of the GFP signal with that of MitoTracker CMTMRos dye confirms the localization of the MPC1 and MPC2 subunits to the mitochondrion. Images were acquired using DeltaVision Deconvolution microscope (GE Healthcare) at 100x magnification and processed using Fiji software (Schindelin et al., 2009). Co-localization of the GFP and MitoTracker signals were examined using the Coloc2 plugin in Fiji and the PCC values for panel A images 1 to 3 are 0.70, 0.84, and 0.84, respectively. The PCC values for panel B images 1 to 3 are 0.97, 0.97, and 0.86, respectively.

### Generation of *Δmpc1, Δmpc2,* marker-excised *Δmpc2* and *Δmpc1Δmpc2* parasites

Next, we decided to examine the essentiality of *mpc1* and *mpc2* genes during the asexual and sexual stages of *P*. *berghei*. The final constructs for knockout of *Pb*MPC1 and *Pb*MPC2 were generated using the recombineering-based method described previously (Pfander et al., 2011) and validated by PCR (Supplementary Figure S5).

*Δmpc1* and *Δmpc2* singly-gene disrupted parasites in *P. berghei* ANKA with C57BL/6 mice as host were generated. Wildtype *P. berghei* ANKA parasites were transfected either with the linearized PbMPC1 or PbMPC2 knockout construct and C57BL/6 mice were infected with the transfected parasites where via double homologous recombination, MPC1 or MPC2 gene are expected to be replaced by the drug selection marker (Figure 4A-C). Drug-resistant transfectants were obtained and genotyping was performed by PCR. 5’ and 3’integration PCRs were carried out to confirm the integration of drug selection marker cassette at the expected locus. Gene-specific forward and reverse primers were used to confirm the absence of *mpc1* gene in *Δmpc1* and for the absence of *mpc2* gene in *Δmpc2* parasites. 5’ and 3’ integration PCRs with gDNA from *Δmpc1 parasites* showed expected bands of 3.1 kb and 8.1 kb, respectively (Supplementary Figure S6C) and likewise, 5’ and 3’integration PCRs with gDNA from *Δmpc2 parasites* showed expected bands of 7.8 kb and 6.6 kb, respectively (Supplementary Figure S6D) whereas PCR with gene-specific primers were not positive for *mpc1* or *mpc2* gene (Supplementary Figure S6E, F). Thus, the generation of *Δmpc1* and *Δmpc2* parasites were successfully achieved. Before phenotyping, limiting dilution cloning of the knockout parasites was performed to obtain clonal parasite lines. The genotype of *Δmpc1* and *Δmpc2* clonal lines were confirmed by PCR (Figure 4D, E). Thereafter, the absence of *Pb*MPC1 gene expression in *Δmpc1* and *Pb*MPC2 gene expression in *Δmpc2* parasites was also validated at the transcript level by RT-PCR (Supplementary Figure S8B, C). The successful generation of individual *Δmpc1* and *Δmpc2* knockout parasites has shown that these genes are not essential during asexual stages of the parasites.

**Figure 4.**
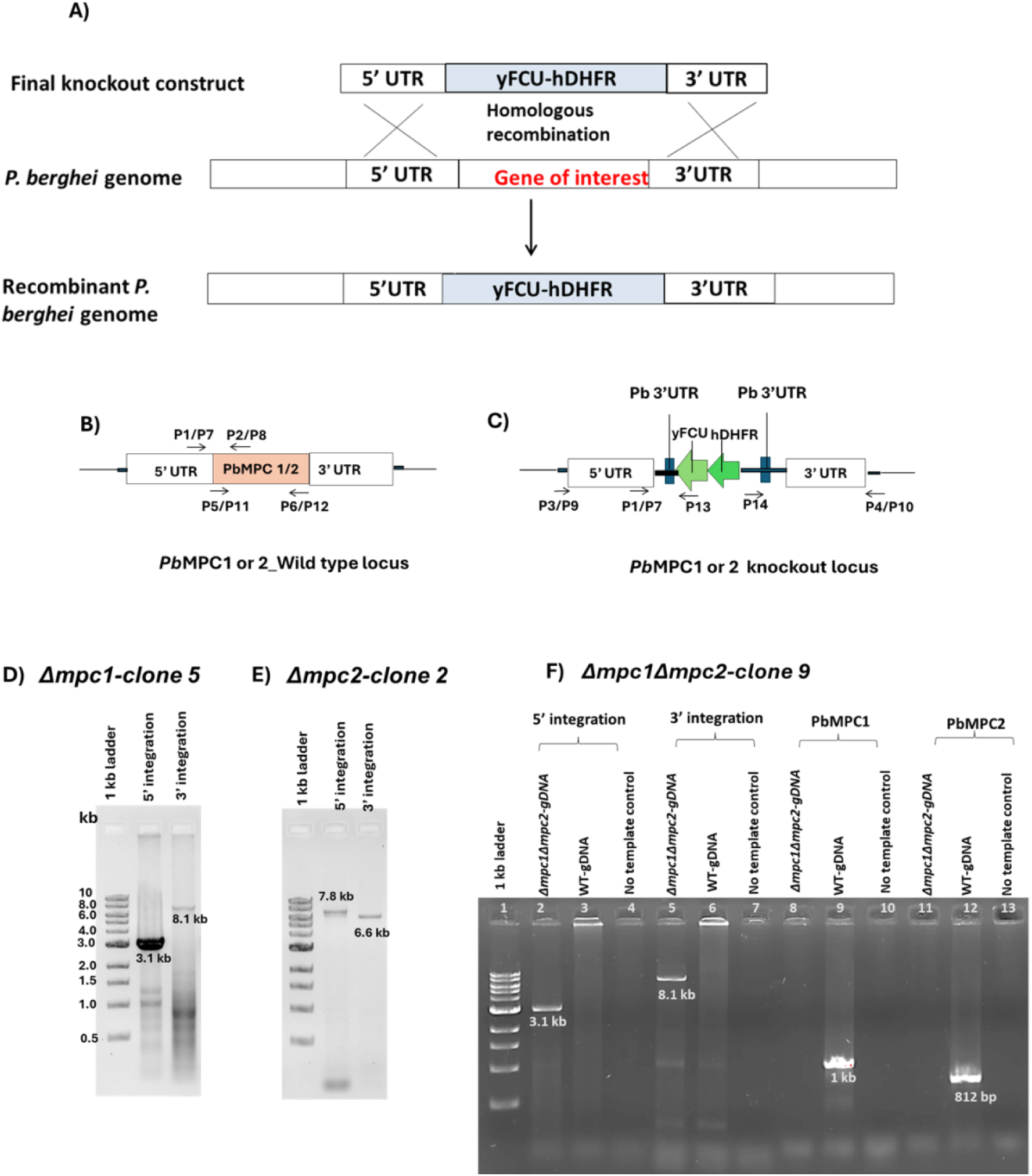
PCR validation of *Δmpc1*, *Δmpc2*, and *Δmpc1Δmpc2* parasite clonal lines. (A) Schematic of the strategy used for the knockout of *mpc1* and *mpc2* genes in *P. berghei* parasites. The constructs for knockout of MPC1 and MPC2 generated by the recombineering method were used for transfection of the wild type *P. berghei ANKA* parasites. Upon transfection with the constructs, through double crossover homologous recombination, the gene of interest was replaced by the selection marker (hDHFR-yFCU), and drug-resistant transfectants were obtained. (B) and (C) Schematics of the MPC1 or 2 gene loci in the genome of *P. berghei* wild-type parasites and of the locus after knockout, respectively. Primers used for PCR genotyping are indicated. D) PCR with gDNA from *Δmpc1* clone 5 parasite line as template and P3/P13 and P4/P14 primers confirmed 5’ and 3’ integration, respectively. E) PCR with gDNA from *Δmpc2* clone 2 parasite line as template and P9/P13 and P10/P14 primers confirmed 5’ and 3’ integration, respectively. Lane 1 in both panels A and B have 1kbp ladder as size marker. F) Confirmation of Δmpc1Δmpc2 parasite clone 9. Lane 1, 1 kb DNA ladder. Lanes 2-4, 5’ integration PCR using P3/P13 primers and gDNA as template (gDNA from *Δmpc1Δmpc2* clone 9 (lane 2), wild-type parasites (lane 3), and without template control (lane 4)). Lanes 5-7, 3’ integration PCRs using P4/P14 primers with gDNA as template (gDNA from *Δmpc1Δmpc2* clone 9 (lane 5), wild-type (lane 6), and without template control (lane 7)). The amplicons of the expected sizes were observed. In *Δmpc1Δmpc2* clone 9, the absence of the PbMPC1(lane 8) and PbMPC2 (lane 11) genes were confirmed using primer pairs; P1-P6 and P7-P8, respectively. Here, PCR with gDNA from wild-type showed expected size amplicons for PbMPC1 (lane 9) and PbMPC2 (lane 12). 1kb to be read as 1kbp (kilo base pairs).

Next, *Δmpc1Δmpc2* parasites were generated to analyse their combined effect on the parasite growth and to compare with *Δmpc1* and *Δmpc2* parasites. Due to the limitation of selection markers for genetic manipulation in *P. berghei*, negative selection on *Δmpc2* gene knockout parasites was carried out to excise the marker (hDHFR-YFCU cassette) by using the toxic metabolite 5-fluorocytosine (5FC) lethal to the parasites (Supplementary Fig.S7). PCR genotyping of 5FC-resistant parasites confirmed the absence of the cassette for the drug-resistant marker in the *Δmpc2* line (Supplementary Figure S7C, D). Thereafter, limiting dilution cloning was performed to isolate marker-excised *Δmpc2* clonal parasite line and this was verified by PCR genotyping prior to use in further studies (Supplementary Figure S7E). The loss of resistance to pyrimethamine was also checked by injecting a marker-excised parasite line into mice and drug pressure was maintained for 25 days. We did not observe the appearance of drug-resistant parasites, confirming the successful excision of the hDHFR-yFCU marker.

For the generation of *Δmpc1Δmpc2* parasites, drug-sensitive *Δmpc2* parasites (clone1) were transfected with linearized PbMPC1 knockout construct and C57BL/6 mice were infected with the transfected parasites. Drug-resistant parasites were obtained, and PCR genotyping yielded expected amplicons of size 3.1 kb for 5’ integration and 8.1 kb for 3’ integration (Supplementary Figure S6F). PCR using gene-specific primers confirmed the absence of both *Pb*MPC1 and *Pb*MPC2 genes (Supplementary Figure S6F) and thus, confirmed the successful generation of *Δmpc1Δmpc2* parasites. Thereafter, limiting dilution cloning was carried out to isolate a clonal line of *Δmpc1Δmpc2* parasites and this was verified by PCR (Figure 4F) and RT-PCR (Supplementary Figure S8D) prior to carrying out phenotypic studies.

### MPC1 and MPC2 subunits are not essential from intraerythrocytic asexual up to the ookinete stages of *P. berghei*

The clonal lines of *Δmpc1*, *Δmpc2,* and *Δmpc1Δmpc2* parasites were examined for their phenotypic traits. The asexual stages of the single and double *mpc* knockout lines proliferated well, within both reticulocytes and normocytes. Further, we did not observe any distinct defect in the cell morphologies of the intra-erythrocytic ring, trophozoite, schizont, and gametocytes stages (Figure 5 and 6). Asexual growth rates of *Δmpc1*, *Δmpc2,* and *Δmpc1Δmpc2* parasites were measured and compared with wildtype parasites, and they did not show any significant differences (Figure 5A, C). This suggests that both genes (*mpc1* and *mpc2*) are not essential during intraerythrocytic stages. Similar to single *mpc1* or *mpc2* knockouts, parasites lacking both genes also did not show any noticeable difference in their asexual growth rates (Figure 5E). The survival rates among mice infected with either wild-type or Δ*mpc1,* Δ*mpc2,* and Δ*mpc1Δmpc2* parasites were not significantly different (Figure 5B, D, F). The enumeration of gametocytes was also carried out and percent of female and male gametocytes were calculated, and they too did not show any significant difference between wildtype and *Δmpc1*, *Δmpc2* and *Δmpc1Δmpc2* knockout parasites (Figure 5G and H). Further, there was no defect in the cellular morphology of the gametocytes (Figure 6B). *In vitro* ookinete conversion rates of wild-type, single *Δmpc1*, Δ*mpc2*, and double *Δmpc1Δmpc2* knockout mutants were also calculated and differences were not found to be significant (Figure 5I-K). There was no well-defined defect observed in morphologies of ookinetes across wildtype, *Δmpc1*, *Δmpc2* and *Δmpc1Δmpc2* parasites (Figure 6C). Overall, these results suggest that the deletion of MPC subunits does not have any impact on parasite morphology, growth and development of the intraerythrocytic stages, gametocyte formation, conversion to gametes, fertilization, zygote formation and conversion to ookinetes.

**Figure 5.**
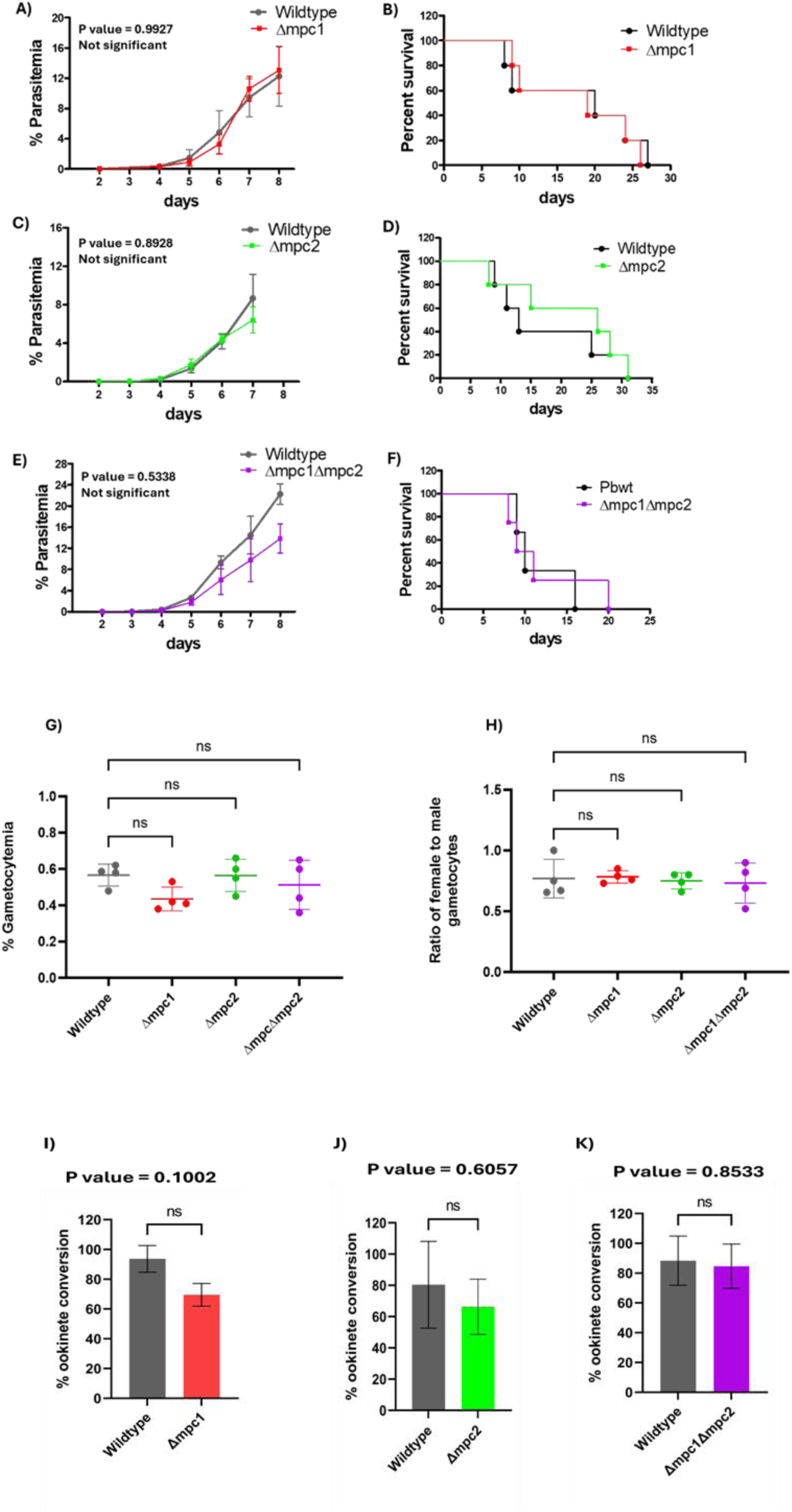
Phenotypic analyses of *Δmpc1, Δmpc2, Δmpc1Δmpc2 P. berghei* lines through intra-erythrocytic asexual, gametocyte, and in vitro generated ookinete stages. (A), (C) and (E) Growth rate of the intra-erythrocytic stages of *Δmpc1, Δmpc2,* and *Δmpc1Δmpc2* parasites in C57BL/6 mice compared with that of the wild type. 10^4^ parasites were injected into mice (N=5 for each genotype) and statistical analysis was performed using Student’s unpaired t-test using GraphPad Prism 9.0. (B), (D) and (F) Percent survival curves of mice infected with either wild-type, *Δmpc1, Δmpc2,* or *Δmpc1Δmpc2* parasites were plotted and calculated using Gehen– Breslow–Wilcoxon test and Log-rank (Mantel-cox) test for Δmpc1 (Log-rank (Mantel-cox) test P value= 0.7679; Gehen–Breslow–Wilcoxon test P value = 0.9176), Δmpc2 (Log-rank (Mantel-cox) test P value= 0.6085; Gehen– Breslow–Wilcoxon test P value = 0.5316), and Δmpc1Δmpc2 parasites (Log-rank (Mantel-cox) test P value= 0.8386; Gehen–Breslow–Wilcoxon test P value = 0.8626). On comparing data obtained with wild-type parasites, no major differences were observed either for asexual growth rates or for survival curves with *mpc* null parasites*. (G-H)* Gametocyte prevalence in wild-type, *Δmpc1, Δmpc2,* and *Δmpc1Δmpc2* parasites. 10^5^ parasites were injected into mice (N=4 for each genotype). Percent gametocytaemia was enumerated for each genotype and statistical analysis was performed. The error bar for each genotype represents mean ± SD from 4 mice. Statistical analysis was performed and a P value of 0.2028 (> 0.05) was obtained using one-way ANOVA test using GraphPad Prism 9.0. (I-K) Enumeration of in vitro ookinete conversion rates of wild-type and *Δmpc1, Δmpc2,* and *Δmpc1Δmpc2* parasites. The data represents mean ± SD values obtained from two mice for each genotype. Statistical analysis was done using the Student’s unpaired t-test using GraphPad Prism 9.0. 10^7^ parasites were injected into mice (N=2 for each genotype) for the experiments in panels (I) and (K) whereas 10^6^ parasites were injected for the experiments with wild-type and Δmpc2 in panel J.

**Figure 6.**
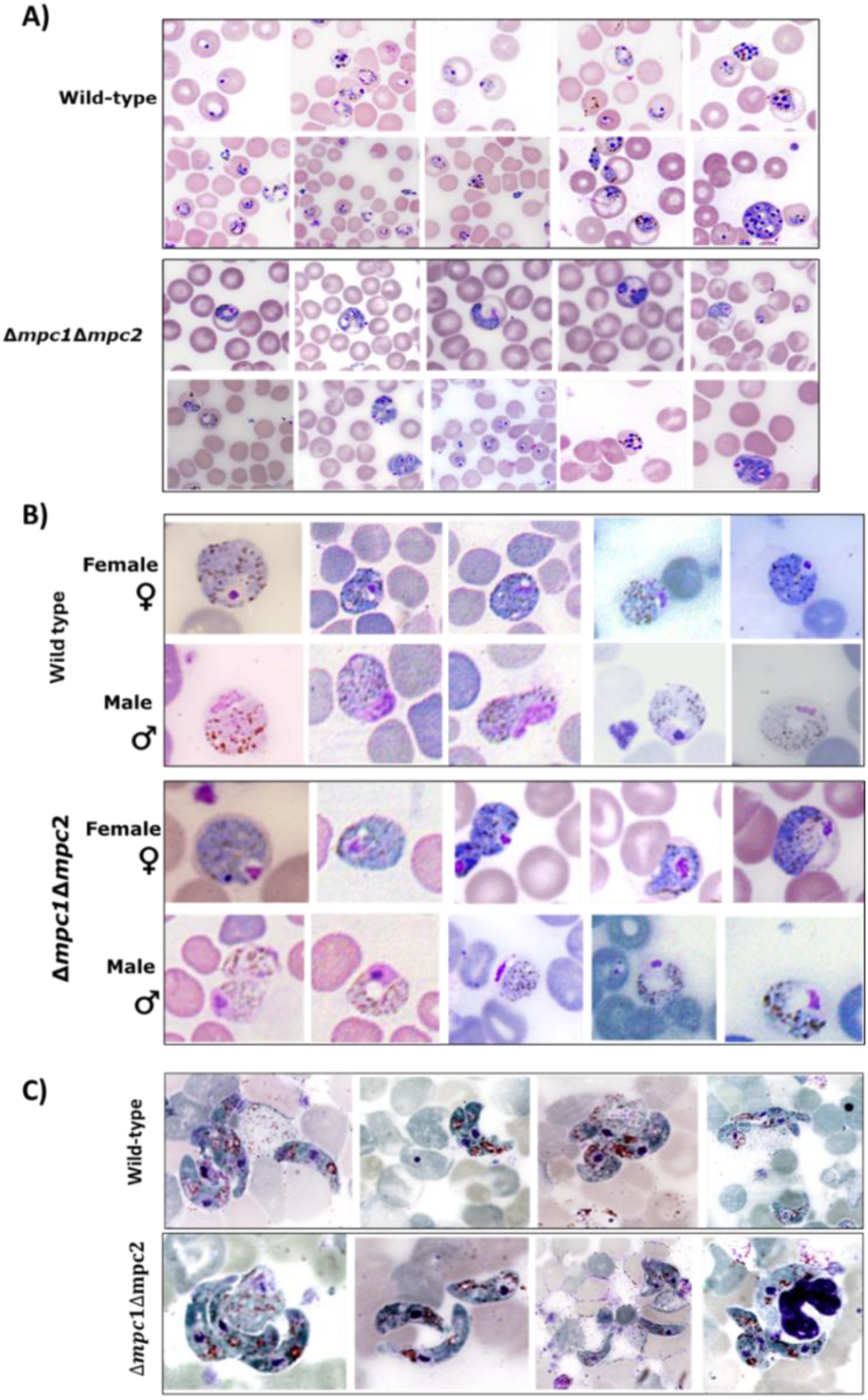
Cell morphologies of *mpc* knockout parasite lines through intra-erythrocytic asexual, gametocyte, and *in vitro* generated ookinete stages. A) Asexual blood stages (ring, trophozoite, and schizont) of wild-type and *Δmpc1Δmpc2* parasites. B) Gametocyte stages. C) Mature ookinete stages. Similar to the wild-type, parasites lacking both the *mpc* genes exhibit normal development within the erythrocytes.

### Isotope tracing LC-MS based metabolomics highlights differences between wild type and Δ*mpc1Δmpc2 P. berghei*

Further investigation was necessary to elucidate why Δ*mpc1Δmpc2* parasites exhibit no significant difference in growth phenotype. To address this, we used isotope tracing LC-MS-based metabolomics to compare the isotope profiles of metabolites from the gametocyte stages of *P. berghei* wild-type and Δ*mpc1Δmpc2* parasites. As the gametocytes of *P. berghei* ANKA could be enriched to > 95 % purity free of asexual stages, all metabolomics studies were carried out with this stage of the parasite. In addition, the male or female gametocytes, that constitute a non-replicative sexual stage of *Plasmodium* sp. rely on the TCA cycle metabolism to generate energy through oxidative phosphorylation (Ke et al., 2015; MacRae et al., 2013, 2012). Relative quantification of the levels of the isotopologues of intracellular metabolites from the central metabolic pathways, such as glycolysis and the TCA cycle was performed on metabolite extracts of *P. berghei* gametocytes fed *in vitro* with either U-^13^C_6_-glucose or U-^13^C ^15^N L-glutamine. Supplementary Figure S9 provides a schematic of the possible isotopologues of the metabolites that can form from labelled glucose and glutamine.

Isotope tracing with U-^13^C_6_-glucose shows decreased levels of M+2 and M+4 isotopologues of TCA <u>cycle intermediates in *Δmpc1Δmpc2 P. berghei* gametocytes.</u> After a 90-minute incubation with U-^13^C_6_-glucose, we detected isotopologues of some of the glycolytic pathway intermediates namely, M+6 isotopologue of glucose-6-phosphate, M+3 (phosphoenol pyruvate) PEP, and M+3 lactate that were at similar levels in both wild-type and *Δmpc1Δmpc2* gametocytes (Figure 8A and Figure 10A). Interestingly, the level of M+3 isotopologue of pyruvate was significantly higher at 60% of the total pool in *Δmpc1Δmpc2* parasites as compared to 24% in the wild-type (Figure 8A). These results suggest a disruption in pyruvate entry into the mitochondrion in the *Δmpc1Δmpc2* parasites, leading to elevated cytosolic pyruvate levels when compared to wild-type parasites. In contrast, lactate levels remained similar in *Δmpc1Δmpc2* and wild-type parasites, likely due to its rapid excretion into the medium.

**Figure 8.**
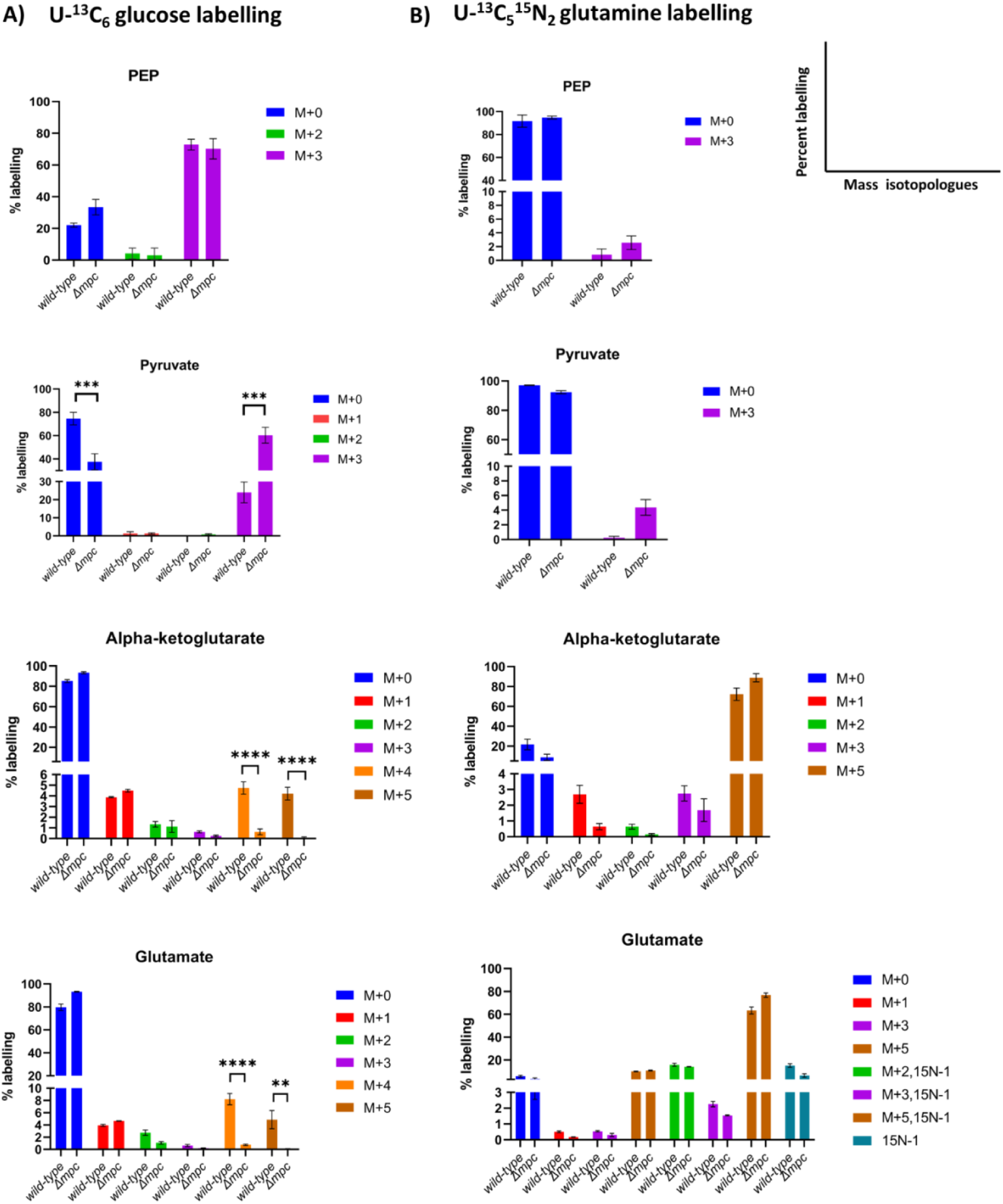
^13^C-enrichment of glycolytic and TCA cycle intermediates extracted from *P. berghei* gametocytes incubated for 90 minutes with U-^13^C_6_-glucose (A) and U-^13^C ^15^N_2_ L-glutamine (B). ^13^C or ^15^N enrichment of a metabolite from wild-type and *Δmpc1Δmpc2* gametocytes are plotted, with the y-axis indicating the percent abundance of each isotopologue of the metabolite relative to the total abundance of all isotopologues of that metabolite. The graphs for percentage labelling of indicated metabolites (PEP, pyruvate, alpha-ketoglutarate, and glutamate) were drawn after correcting for natural abundance (NA) using IsocorrectoR. The labelling data from wild-type and *Δmpc1Δmpc2* gametocytes represent N= 2 independent biological replicates, with error bars indicating SD. Significance determined by unpaired t-test is highlighted in the figure, with p-values of ≤0.05, ≤0.01, ≤0.001, and ≤0.0001 indicated by one, two, three, and four asterisks, respectively.

Moving from glycolytic intermediates into the TCA cycle, consistent detection of M+2 isotopologue of acetyl-CoA in all ^13^C_6_-glucose-labelled samples proved challenging precluding comparative analysis of acetyl-CoA levels between the wild-type and *Δmpc1Δmpc2* gametocytes. Surprisingly, both M+2 and M+4 isotopologues of citrate could be detected in *Δmpc1Δmpc2* parasites indicating a flux of two rounds through the cycle despite the absence of the transporter. However, the extracted ion chromatograms of citrate were often very broad hindering reliable quantification. In contrast, the M+4 isotopologues of succinate, fumarate, and malate could be consistently detected and reliably quantified across wild type and the knockout parasites. The presence of M+4 isotopologues of succinate, fumarate and malate further supported the presence of labelled pyruvate in the mitochondrion and its subsequent utilization in the TCA cycle. However, the *Δmpc1Δmpc2* gametocytes exhibited a significant decrease in the levels of the M+4 isotopologues of these metabolites, (succinate (24-fold), fumarate (30-fold), and malate (35-fold)) suggesting the presence of lower levels of labelled pyruvate in the mitochondrion (Figure 9A). Our study also detected M+2 and M+4 isotopologues of aspartate in both parasite types with the latter at 14-fold lower levels in the knockout gametocytes (Figure 9A). The presence of M+2 and M+4 aspartate indicate its generation through the TCA cycle flux. Our detection of M+2 and M+4 isotopologues of succinate, fumarate, malate, and aspartate indicates the continued presence of labelled pyruvate albiet at lower levels in the mitochondrion, highlighting that mpc is a key transporter of pyruvate. It should be noted that during the incubation of parasites with U-^13^C_6_-glucose, glutamine is available and would serve as an alternate carbon source to fuel the TCA cycle.

**Figure 9.**
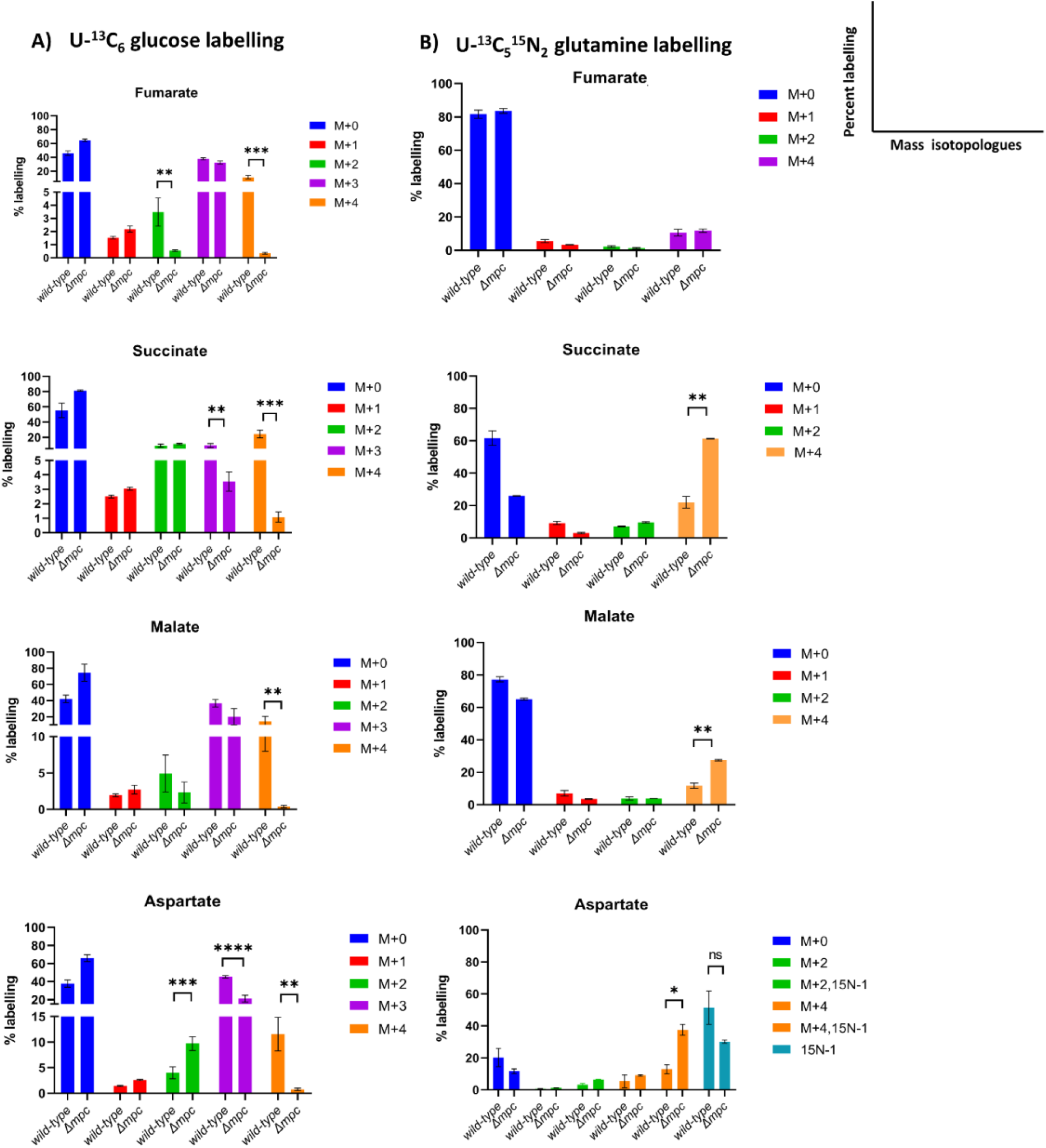
^13^C-enrichment of TCA cycle intermediates from *P. berghei* gametocytes incubated for 90 minutes with U-^13^C_6_-glucose (A) and U-^13^C_5_^15^N_2_ L-glutamine (B). ^13^C or ^15^N enrichment of a metabolite from wild-type and *Δmpc1Δmpc2* gametocytes are plotted, with the y-axis indicating the percent abundance of each isotopologue of the metabolite relative to the total abundance of all isotopologues of that metabolite. The graphs for percentage labelling of indicated metabolites (fumarate, succinate, malate and aspartate) were plotted after correcting for natural abundance (NA) using IsocorrectoR. Error bar shows SD of N=2 independent biological replicates. Significance determined by unpaired t-test is highlighted in the figure, with p-values of ≤0.05, ≤0.01, ≤0.001, and ≤0.0001 indicated by one, two, three, and four asterisks, respectively.

In addition to detecting M+2 and M+4 isotopologues of the TCA cycle intermediates fumarate, malate, alpha-ketoglutarate, and succinate, and through oxaloacetate (OAA), aspartate arising from the first and second turn of the cycle, we also observed their M+3 and M+5 isotopologues. The M+3 isotopologue arises from two pathways. First, through anaplerotic reactions that replenish the TCA cycle intermediates. Within the parasite cytosol, the M+3 isotopologue of OAA produced from M+3 PEP is converted to the M+3 malate by cytosolic malate dehydrogenase and the M+3 isotopologue of fumarate is produced as a by-product of AMP synthesis. Both M+3 malate and M+3 fumarate enter the mitochondrion and drive the TCA cycle (Supplementary Figure S9). In the second scenario, the M+3 isotopologues of the TCA cycle intermediates; alpha-ketoglutarate, fumarate, succinate, malate, OAA, and aspartate can also be obtained via 2^nd^ turn of the TCA cycle (Buescher et al., 2015).

The M+4 isotopologue of citrate generated during the second round produces alpha-ketoglutarate with two isotopologue, M+4 and M+3, populations. These two populations arise due to differences in the labelling pattern of citrate (a six-carbon metabolite) with four ^13^C-carbon atoms (Buescher et al., 2015). Subsequently, M+3 succinate, malate, fumarate, and aspartate can also be formed. In our study, no significant differences were observed for M+3 isotopologues of fumarate in both the parasite types. The levels of M+3 isotopologues of malate, succinate, and aspartate were slightly lower (1.5-fold for malate; 2.5-fold for succinate, and 2-fold for aspartate) in the *Δmpc1Δmpc2* gametocytes compared to wild-type parasites (Figure 9A).

The detection of M+5 isotopologue of citrate, in *Δmpc1Δmpc2* gametocytes, indicates flux through M+3 isotopologue of OAA and M+2 isotopologue of acetyl-CoA. The M+3 OAA can be generated in the mitochondrion directly from the M+3 isotopologue of malate or from the M+3 isotopologue of fumarate that enter the mitochondrion and feed into the TCA cycle anaplerotically (Supplementary Figure S9). Earlier studies have shown the anaplerotic entry of fumarate into the TCA cycle (Bulusu et al., 2011; Ke et al., 2015; Srivastava et al., 2016). Further, M+5 isotopologues of alpha-ketoglutarate and glutamate were identified in wild-type and *Δmpc1Δmpc2* parasites. Although an extremely low percentage (4 %) of M+5 isotopologues of alpha-ketoglutarate and glutamate was found in the wild-type, this percentage dropped to about 0.1 % in *Δmpc1Δmpc2* parasites (Figure 8A). Interestingly, the M+3-isotopologue of citrate, which could have originated from the M+3 OAA and unlabelled acetyl-CoA, was absent in both wild-type and *Δmpc1Δmpc2* gametocytes. This absence indicates the presence of sufficient pools of the M+2 isotopologue of acetyl-CoA produced from U-^13^C_6_-glucose-derived pyruvate and transport of unlabelled acetyl-CoA, if any from the cytosol must be very minor. However, it should be noted that the extracted ion chromatogram of citrate was broad, possibly due to matrix effects. In summary, the M+5-isotopologue of citrate was detected and not the M+3-isotopologue of citrate.

These outcomes suggest that *P. berghei* MPC plays a role in the transport of pyruvate for its subsequent conversion to acetyl-CoA. The maintenance of the acetyl-CoA pool is crucial for the completion of the TCA cycle. The identification of M+2 and M+4 isotopologues of citrate in *Δmpc1Δmpc2* gametocytes indicates that despite the deletion of the two *mpc* genes, the M+2 isotopologue of acetyl-CoA is produced at a moderate rate in the mitochondrial matrix.

An interesting finding of this U-^13^C_6_-glucose labelling study was the detection of M+3 isotopologue of alanine in both *Δmpc1Δmpc2* and wild-type gametocytes at levels of 35 % in the wild-type and 50 % in *Δmpc1Δmpc2* parasites (Figure 10A). This suggests transamination of pyruvate which may be occurring in the cytosol or the mitochondrion or both compartments.

**Figure 10.**
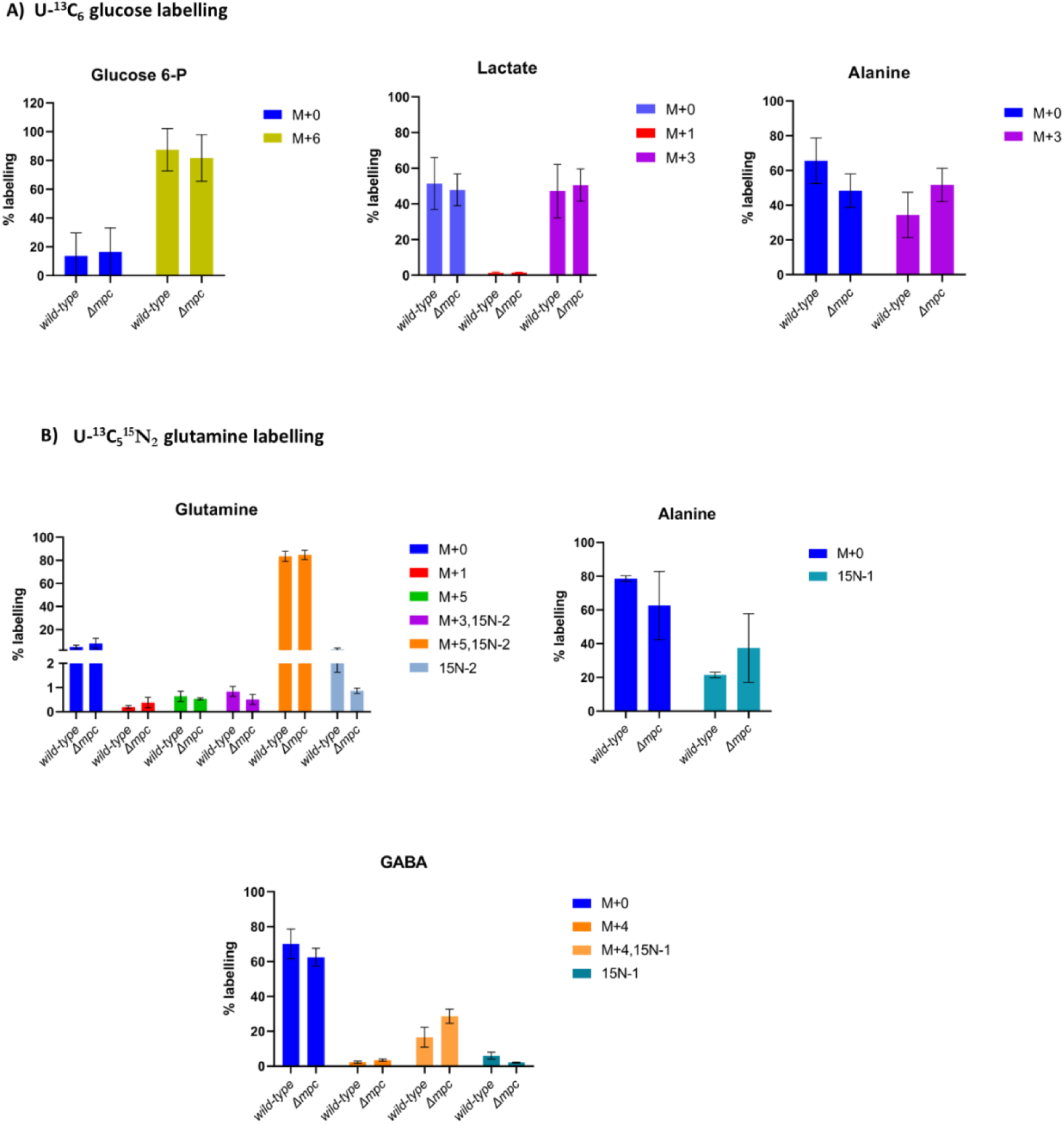
^13^C-enrichment of glycolytic and TCA cycle intermediates from *P. berghei* gametocytes incubated for 90 minutes with U-^13^C_6_-glucose (A) and U-^13^C ^15^N_2_ L-glutamine (B). ^13^C or ^15^N enrichment of a metabolite from wild-type and *Δmpc1Δmpc2* gametocytes are plotted, with the y-axis indicating the percent abundance of each isotopologue of the metabolite relative to the total abundance of all isotopologues of that metabolite. The graphs for percentage labelling of indicated metabolites (glucose-6-phosphate, lactate, alanine, glutamine, and GABA (Gamma-aminobutyric acid), were plotted after correcting for natural abundance (NA) using IsocorrectoR. Error bar shows SD of N=2 independent biological replicates. Here, the significance determined by the unpaired t-test is highlighted in the figure, with p-values of ≤0.05.

Isotope tracing with U-^13^C ^15^N L-glutamine shows a minor difference in flux through TCA cycle in <u>Δ*mpc1Δmpc2 P. berghei* gametocytes.</u> Both parasite types exhibited the presence of M+5 isotopologues of glutamate and alpha-ketoglutarate at similar levels. The M+4 isotopologues of succinate and malate showed a minor increase of 2.7- and 2.4-fold, respectively in *Δmpc1Δmpc2 P. berghei* gametocytes (Figure 8B, 9B). Similarly, the level of M+4-^15^N_1_ isotopologue of aspartate was 3-fold higher (Figure 9B). These observations from U-^13^C ^15^N_2_ L-glutamine labelling studies suggest a minor increase in glutamine utilisation in *Δmpc1Δmpc2* gametocytes, possibly necessary for maintaining optimum flux through the TCA cycle in the absence of the pyruvate transporter.

We also observed M+3 isotopologues of PEP and pyruvate, metabolites of the glycolytic pathway, albeit at a very low percentage (<4 %) of the total (Figure 8B). This labelled PEP is generated from labelled OAA by the enzyme, phosphoenolpyruvate carboxykinase, followed by conversion to labelled pyruvate (Supplementary Figure S9B). The detection of both M+2 and M+4 isotopologues of citrate in *Δmpc1Δmpc2* from U-^13^C_5_^15^N_2_ L-glutamine-labelling studies provides additional confirmation of the presence of acetyl-CoA pool within the mitochondrion. M+4 citrate is derived from the condensation of M+4 isotopologue of OAA and acetyl-CoA containing two unlabelled acetyl carbons, while M+2 labelled citrate is derived from M+2 isotopologue of acetyl-CoA and unlabelled OAA. However, relative quantification of citrate between the two parasite lines was not possible due to its broad extracted ion chromatogram. The levels of GABA derived from U-^13^C ^15^N L-glutamine remain unchanged across wild-type and *Δmpc1Δmpc2* gametocytes (Figure 10B).

In both *Δmpc1Δmpc2* and wild-type *P. berghei* gametocytes, the detection of ^15^N-labeled alanine at similar levels (Figure 10B) provides further evidence for pyruvate-to-alanine interconversion mediated by a transaminase. It is possible that ^13^C_5_^15^N_1_-glutamate or ^13^C_4_^15^N_1_-aspartate serves as an amino group donor, transferring their ^15^N-atom to pyruvate, resulting in ^15^N isotopologue of alanine in addition to the formation of ^13^C_5_-alpha-ketoglutarate or ^13^C_4_-OAA, respectively. Another potential donor for the ^15^N amino group could be GABA, which undergoes conversion to succinic semialdehyde through a putative mitochondrial transaminase (PBANKA_010740, OAT), concurrently producing alanine from pyruvate.

## Discussion

Pyruvate, a central metabolite produced through the glycolytic pathway, regulates carbohydrate, fatty acid, and amino acid metabolisms. The entry of pyruvate into mitochondria is enabled by MPC comprising two subunits, MPC1 and MPC2. We have shown the localization of MPC1 and MPC2 to the mitochondrion in *P. berghei*. Our results from a functional complementation assay confirm that both subunits of *Plasmodium* MPC are necessary for pyruvate transport activity and individual subunits of MPC (MPC1 or MPC2) cannot form a functional homo-complex that mediates the uptake of pyruvate. In glucose labelling experiments, an increase in levels of M+3 pyruvate in Δ*mpc1*Δ*mpc2 P. berghei* clearly suggests the role of MPC in pyruvate transport. Though M+4 succinate, fumarate and malate could be detected, their levels are drastically reduced (14-35-fold) in Δ*mpc1*Δ*mpc2 P. berghei* again indicating that pyruvate transport into the mitochondrion is dramatically impaired. Overall, these results indicate that MPC is the major carrier albeit with alternate routes of entry being present. It should be noted that we are calculating relative amounts of individual isotopologues of a metabolite, and this does not factor in anaplerotic fluxes namely M+3 fumarate and malate. In addition, alpha-ketoglutarate entry through glutamine anaplerosis does also occur as seen from our ^13^C_5_^15^N_2_ glutamine labelling studies and also by Srivastava and co-workers (Srivastava et al., 2016). In *T.brucei* too, knockout of MPC leads to the accumulation of pyruvate (Štáfková et al., 2016).

Recently, using CRISPR technology, Prigge and coworkers have successfully generated P*. falciparum* lines lacking both *mpc1* and *mpc2* genes and these showed only a minor growth defect. Acetyl-CoA pools derived from ^13^C_6_-glucose were significantly reduced in Δ*mpc1*Δ*mpc2 P. falciparum* but their continued presence suggested alternate carriers operating in the parasite (Rajaram et al., 2024). In our study, Δ*mpc1,* Δ*mpc2,* and Δ*mpc1*Δ*mpc2 P. berghei* showed no defect through different developmental stages *viz.,* asexual blood stages, gametocytes and *in-vitro* developed ookinete stages. Support for continued pyruvate entry into the mitochondria of the Δ*mpc1*Δ*mpc2* of *P. berghei* comes from continued presence of equal levels of M+2 succinate and increased levels of M+2 aspartate from glucose labelling when compared with the wild type. Simultaneous knockout of both BCKDH and KDH in *P. falciparum* was possible only in the presence of exogenously added acetate indicating that the mitochondrial acetyl-CoA pool is essential for the parasite (Nair et al., 2023; Oppenheim et al., 2014). The absence of growth defects in *P. berghei* up to ookinete stages despite the lack of MPC has led us to examine 3 possibilities in Δ*mpc* parasites that enable their survival.

First, the existence of an alternate mitochondrial carrier with substrate specificity for pyruvate. Prigge and co-workers (Rajaram et al., 2024) in their recent study have shown that simultaneous knockout of *mpc1*, *mpc2*, dicarboxylate/tricarboxylate carrier (*dtc*) and oxoglutarate carrier (*ogc*) genes did not further affect incorporation of glucose derived carbon into acetyl-CoA over what was observed for Δ*mpc1*Δ*mpc2* establishing that OGC and DTC do not transport pyruvate. Currently, we do not know if any of the other transporter/s could transport pyruvate into the mitochondrion.

The second possibility in Δ*mpc* parasites could be the entry of acetyl-CoA from the cytosol into the mitochondrion that would lead to the restoration of mitochondrial acetyl-CoA and subsequently citrate pools. To address the possible transport of acetyl-CoA from the cytosol into the mitochondrion, contributing to the citrate pool in Δ*mpc1*Δ*mpc2 P. berghei*, we examined for the presence of M+3 isotopologue of citrate in extracts of the knockout parasites fed with ^13^C_6_-glucose. The M+3 isotopologue of citrate can be derived from M+3 OAA (generated by MQO) condensing with unlabelled acetyl-CoA imported from the cytosol into the mitochondrion. We did not observe the M+3 isotopologue of citrate in both wild-type and Δ*mpc1*Δ*mpc2 P. berghei*. However, we detected M+2 citrate (arising from unlabelled OAA and M+2 acetyl-CoA), and M+5 citrate (arising from M+3 OAA and M+2 acetyl-CoA) in both parasite lines. Recent studies on *P. falciparum* lines lacking the E2 subunit of *Pf*BCKDH and *Pf*KDH or ΔLipL2 genes led to a complete block of mitochondrial acetyl-CoA pools and the parasites survived only when the medium was supplemented with excess acetate. Under these conditions, from ^13^C_6_-glucose labelling experiments, the authors provide evidence of acetyl-CoA transport from the cytosol into the mitochondrion (Nair et al., 2023). As the sensitivity of the LC-MS technique is very high, our inability to detect M+3 citrate in both wild-type and Δ*mpc1*Δ*mpc2* parasites indicates either the complete absence of this flux or it being extremely minor in *P. berghei*.

Third, is the existence of an alternate route for the generation of pyruvate within the mitochondrion leading to the observed normal growth phenotype despite lacking MPC. In mammalian cells, pyruvate produced in the cytosol can be converted to alanine by the cytosolic enzyme, L-alanine aminotransferase 1 (ALT1; also known as GPT1). Alanine thus generated is transported into the mitochondria and is converted to pyruvate by the mitochondrial L-alanine aminotransferase 2 (ALT2; also known as GPT2). In *T. gondii,* upon the deletion of *mpc1* gene, parasites showed a minor defect in the growth and development, thus alternate routes of pyruvate or acetyl-CoA generation were suggested within the mitochondrion (Lyu et al., 2023). Here, the gene encoding the ALT enzyme is annotated although the biochemical function and localization are yet to be established. However, the *Plasmodium* genome lacks a gene similar to known ALT1 or ALT2.

In both ^13^C_6_-glucose and ^13^C_5_^15^N_2_-glutamine labelling experiments, we consistently detected significant levels of isotope-labelled alanine in both Δ*mpc1*Δ*mpc2* and wild type *P. berghei* while in extracts of uninfected erythrocytes, the levels of M+3 and ^15^N_1_ isotopologues of alanine were very low (7 % of M+3 alanine from ^13^C_6_-glucose and only 1% of ^15^N_1_ alanine from ^13^C_5_^15^N_2_-glutamine) (Supplementary Figure S10). As the erythrocyte contamination in our *P. berghei* gametocytes samples was less than 10 %, the main contributor to the isotopically labelled alanine is the parasite. These results suggest that pyruvate can undergo a transamination reaction to form alanine in both *P. berghei* lines. Transport of alanine into the mitochondrion followed by conversion back to pyruvate within the organelle would serve as an alternate pathway for the presence of pyruvate even in the absence of MPC. Hence, a mitochondrial transamination reaction could be a potential reason why the growth rate of Δ*mpc1*Δ*mpc2 P. berghei* did not differ substantially from the wild-type. These results from Δ*mpc1*Δ*mpc2 P. berghei* can be correlated to a previous study on *Arabidopsis thaliana* by Le and co-workers (Le XH et al., 2021) wherein the authors showed that plant growth was not affected by the loss of MPC genes whereas an extremely retarded phenotype was obtained when the ALT gene was deleted in the background of *mpc* knockout. The study on *mpc* knockout *Arabidopsis* highlighted the contribution of mitochondrial transamination reaction mediated by ALT in supplying pyruvate from alanine for the maintenance of the TCA cycle required to sustain plant growth. The proposed pathway for the alternate supply of pyruvate from alanine within the mitochondrion of *P. berghei* is shown in Figure 11.

**Figure 11.**
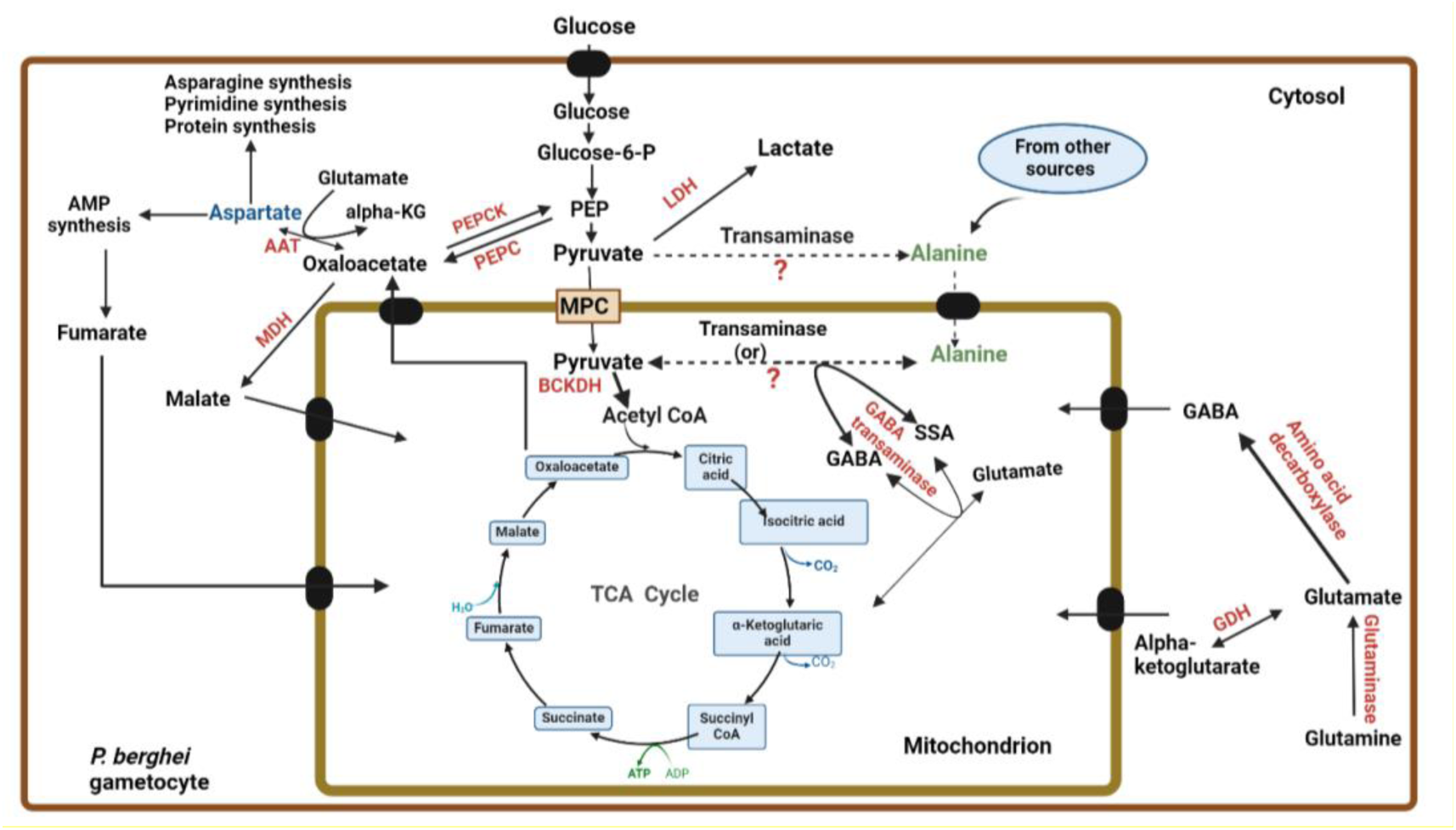
An alternate pathway for the generation of pyruvate in the mitochondrion of *P. berghei*. From the results of our metabolomics study on Δ*mpc1*Δ*mpc2* and wild-type parasites, we propose that the mitochondrial pyruvate pool is maintained by two routes; 1) MPC and 2) a combination of an alanine transporter enabling mitochondrial uptake of the amino acid along with a mitochondrial transaminase that converts alanine to pyruvate within the organelle. Based on our results obtained from ^13^C_6_-glucose and ^13^C_5_^15^N_2_ L-glutamine labelling experiments, we propose transamination reactions occurring in both the cytosol (by a transaminase that converts pyruvate to alanine) and in the mitochondrion (by a transaminase converting alanine to pyruvate). The detection of labelled alanine from both ^13^C_6_-glucose and ^13^C ^15^N L-glutamine studies indicates that a transaminase that converts pyruvate to alanine is present and the potential donor of the amino group could be glutamate. The impact of the loss of the MPC genes can be compensated by a mitochondrial transamination reaction.

Most aminotransferase or transaminase reactions are reversible and require pyridoxal-5’ phosphate as a coenzyme (Kronenberger et al., 2014) in catalysing the reaction between the amino donor and keto acid acceptor (Koper et al., 2023; Kronenberger et al., 2014). A keyword search in PlasmoDB yielded two aminotransferases, aspartate aminotransferase (AAT) (PBANKA_0302300) and a putative GABA transaminase/ornithine aminotransferase (PBANKA_0107400) in the *P. berghei* genome. *Pf*AAT is present in the cytosol whereas, the intracellular localization of a GABA transaminase/ornithine aminotransferase has not yet been experimentally validated (Jortzik et al., 2010; Wrenger et al., 2011). The gene encoding AAT in *P. berghei* is essential for the parasite (Srivastava et al., 2015). Although, the protein sequence of *Plasmodium* AAT exhibits significant variation from both eukaryotic and bacterial counterparts (Wrenger et al., 2011) its activity on pyruvate has not been detected.

In our study, consistent levels of isotopically labelled GABA were detected in both parasite lines. The *P. berghei* genome encodes a gene required for GABA synthesis (PBANKA_1003400; amino acid decarboxylase), GABA transport (PBANKA_0306700), and possibly GABA transamination (PBANKA_0107400; ornithine aminotransferase). Unlike *T. gondii, Plasmodium* lacks a canonical GABA shunt since it lacks a homolog of succinate semialdehyde dehydrogenase required for the generation of the TCA cycle intermediate, succinate from succinic semialdehyde (MacRae et al., 2013, 2012; Srivastava et al., 2016). In a previous study on *P. falciparum* gametocytes, the detection of labelled alanine from both ^13^C_6_-glucose and ^13^C_5_-glutamine labelling suggested the role of GABA in mitochondrial transamination reactions (MacRae et al., 2013, 2012). However, in *Plasmodium*, the role of GABA in regulating the TCA cycle flux through transamination reactions remains poorly understood.

Taken together, our studies on Δ*mpc* parasites suggest a non-essential role for MPC in *P. berghei* up to the ookinete stages. Whether Δ*mpc* parasites exhibit growth defects in other mosquito stages, such as in oocyst maturation, sporozoite development, and hepatocyte stages, remains to be studied. This study also sheds light on the possible existence of mitochondrial transamination reactions that play a role in generating pyruvate from alanine within the mitochondrion.

## Supporting information

Supple_MPC_bioRxiv

## Acknowledgements

HB acknowledges funding from Science and Engineering Research Board (SERB), Department of Science and Technology, Government of India (Grant Number: CRG/2019/004150), JC BOSE fellowship, Science and Engineering Research Board (SERB), Department of Science and Technology, Government of India (Grant number: JCB/2019/000006), Department of Biotechnology, Government of India (Grant number: BT/INF/22/SP27679/2018, Life science research, education and training at JNCASR) and intramural funding from JNCASR. AB acknowledges Counsil of Scientific and Industrial Research (CSIR), Government of India for junior and senior research fellowships. Authors acknowledge the mass spectrometry facility at the Molecular Biology and Genetic Unit (MBGU), JNCASR for all mass spectral data and the animal house facility, JNCASR for supply of mice used in the study. Authors thank Prof. Ravi Manjithaya, MBGU, JNCASR for providing access to the Delta Vision microscope. Authors also thank Anusha Chandrashekarmath for optimising conditions for LC-MS based metabolomics and subsequent data analysis involving corrections for natural isotope abundance and use of El-MAVEN.

## REFERENCES

Agrawal, S., Kumar, S., Sehgal, R., George, S., Gupta, R., Poddar, S., Jha, A., & Pathak, S. (2019). El-MAVEN: A Fast, Robust, and User-Friendly Mass Spectrometry Data Processing Engine for Metabolomics. In A. D’Alessandro (Ed.), High-Throughput Metabolomics: Methods and Protocols (pp. 301–321). Springer New York. 10.1007/978-1-4939-9236-2_19

Aurrecoechea, C., Brestelli, J., Brunk, B.P., Dommer, J., Fischer, S., Gajria, B., Gao, X., Gingle, A., Grant, G., Harb, O.S., Heiges, M., Innamorato, F., Iodice, J., Kissinger, J.C., Kraemer, E., Li, W., Miller, J.A., Nayak, V., Pennington, C., Pinney, D.F., Roos, D.S., Ross, C., Stoeckert, C.J., Treatman, C., Wang, H., 2009. PlasmoDB: A functional genomic database for malaria parasites. Nucleic Acids Res. 37, 539–543. 10.1093/nar/gkn814

Bender, T., Martinou, J., 2016. Biochimica et Biophysica Acta The mitochondrial pyruvate carrier in health and disease: To carry or not to carry? ☆. BBA - Molecular Cell Research 1863, 2436–2442. 10.1016/j.bbamcr.2016.01.017

Bricker, D.K., Taylor, E.B., Schell, J.C., Orsak, T., Boutron, A., Chen, Y.C., Cox, J.E., Cardon, C.M., Van Vranken, J.G., Dephoure, N., Redin, C., Boudina, S., Gygi, S.P., Brivet, M., Thummel, C.S., Rutter, J., 2012. A mitochondrial pyruvate carrier required for pyruvate uptake in yeast, Drosophila, and humans. Science (1979) 336, 96–100. 10.1126/science.1218099

Buescher, J.M., Antoniewicz, M.R., Boros, L.G., Burgess, S.C., Brunengraber, H., Clish, C.B., DeBerardinis, R.J., Feron, O., Frezza, C., Ghesquiere, B., Gottlieb, E., Hiller, K., Jones, R.G., Kamphorst, J.J., Kibbey, R.G., Kimmelman, A.C., Locasale, J.W., Lunt, S.Y., Maddocks, O.D.K., Malloy, C., Metallo, C.M., Meuillet, E.J., Munger, J., Nöh, K., Rabinowitz, J.D., Ralser, M., Sauer, U., Stephanopoulos, G., St-Pierre, J., Tennant, D.A., Wittmann, C., Vander Heiden, M.G., Vazquez, A., Vousden, K., Young, J.D., Zamboni, N., Fendt, S.M., 2015. A roadmap for interpreting 13C metabolite labeling patterns from cells. Curr Opin Biotechnol 34, 189–201. 10.1016/j.copbio.2015.02.003

Bulusu, V., Jayaraman, V., Balaram, H., 2011. Metabolic fate of fumarate, a side product of the purine salvage pathway in the intraerythrocytic stages of Plasmodium falciparum. Journal of Biological Chemistry 286, 9236–9245. 10.1074/jbc.M110.173328

Chai, Y., Wang, C., Liu, W., Fan, Y., Zhang, Y., 2019. MPC1 deletion is associated with poor prognosis and temozolomide resistance in glioblastoma. J Neurooncol. 10.1007/s11060-019-03226-8

Chambers, M.C., MacLean, B., Burke, R., Amodei, D., Ruderman, D.L., Neumann, S., Gatto, L., Fischer, B., Pratt, B., Egertson, J., Hoff, K., Kessner, D., Tasman, N., Shulman, N., Frewen, B., Baker, T.A., Brusniak, M.Y., Paulse, C., Creasy, D., Flashner, L., Kani, K., Moulding, C., Seymour, S.L., Nuwaysir, L.M., Lefebvre, B., Kuhlmann, F., Roark, J., Rainer, P., Detlev, S., Hemenway, T., Huhmer, A., Langridge, J., Connolly, B., Chadick, T., Holly, K., Eckels, J., Deutsch, E.W., Moritz, R.L., Katz, J.E., Agus, D.B., MacCoss, M., Tabb, D.L., Mallick, P., 2012. A cross-platform toolkit for mass spectrometry and proteomics. Nat Biotechnol 30, 918–920. 10.1038/nbt.2377

Chan, X.W.A., Wrenger, C., Stahl, K., Bergmann, B., Winterberg, M., Müller, I.B., Saliba, K.J., 2013. Chemical and genetic validation of thiamine utilization as an antimalarial drug target. Nature Communications 2013 4:14, 1–11. 10.1038/ncomms3060

Creek, D.J., Jankevics, A., Breitling, R., Watson, D.G., Barrett, M.P., Burgess, K.E.V., 2011. Toward global metabolomics analysis with hydrophilic interaction liquid chromatography-mass spectrometry: Improved metabolite identification by retention time prediction. Anal Chem 83, 8703–8710. https://pubs.acs.org/doi/full/10.1021/ac2021823

Daniel Gietz, R., Woods, R.A., 2002. Transformation of yeast by lithium acetate/single-stranded carrier DNA/polyethylene glycol method, in: Guthrie, C., Fink, G.R. (Eds.), Guide to Yeast Genetics and Molecular and Cell Biology - Part B, Methods in Enzymology. Academic Press, pp. 87–96. 10.1016/S0076-6879(02)50957-5

Ghosh, A., Tyson, T., George, S., Hildebrandt, E.N., Steiner, J.A., Madaj, Z., Schulz, E., Machiela, E., Mcdonald, W.G., Galvis, M.L.E., Kordower, J.H., Raamsdonk, J.M. Van, Colca, J.R., Brundin, P., 2016. Mitochondrial pyruvate carrier regulates autophagy, inflammation, and neurodegeneration in experimental models of Parkinson’s disease Sci. Transl. Med. 8, 368ra174. 10.1126/scitranslmed.aag2210.

Divakaruni, A.S., Wallace, M., Buren, C., Martyniuk, K., Andreyev, A.Y., Li, E., Fields, J.A., Cordes, T., Reynolds, I.J., Bloodgood, B.L., Raymond, L.A., Metallo, C.M., Murphy, A.N., 2017. Inhibition of the mitochondrial pyruvate carrier protects from excitotoxic neuronal death. J Cell Biol. 216, 1091–1105. 10.1083/JCB.201612067

Foth, B.J., Stimmler, L.M., Handman, E., Crabb, B.S., Hodder, A.N., McFadden, G.I., 2005. The malaria parasite Plasmodium falciparum has only one pyruvate dehydrogenase complex, which is located in the apicoplast. Mol. Microbiol. 55, 39–53. 10.1111/J.1365-2958.2004.04407.X

Gascuel, O., 1997. BIONJ: An improved version of the NJ algorithm based on a simple model of sequence data. Mol Biol Evol 14, 685–695. 10.1093/oxfordjournals.molbev.a025808

Giannangelo, C., Siddiqui, G., de Paoli, A., Anderson, B.M., Edgington-Mitchell, L.E., Charman, S.A., Creek, D.J., 2020. System-wide biochemical analysis reveals ozonide antimalarials initially act by disrupting Plasmodium falciparum haemoglobin digestion, PLoS Pathog. 16, e1008485. 10.1371/journal.ppat.1008485

Gomes, A.R., Bushell, E., Schwach, F., Girling, G., Anar, B., Quail, M.A., Herd, C., Pfander, C., Modrzynska, K., Rayner, J.C., Billker, O., 2015. A genome-scale vector resource enables high-throughput reverse genetic screening in a malaria parasite. Cell Host Microbe 17, 404–413. 10.1016/j.chom.2015.01.014

Heinrich, P., Kohler, C., Ellmann, L., Kuerner, P., Spang, R., Oefner, P.J., Dettmer, K., 2018. Correcting for natural isotope abundance and tracer impurity in MS-, MS/MS- and high-resolution-multiple-tracer-data from stable isotope labeling experiments with IsoCorrectoR. Sci Rep 8, 1–10. 10.1038/s41598-018-36293-4

Herzig, S., Raemy, E., Montessuit, S., Veuthey, J.L., Zamboni, N., Westermann, B., Kunji, E.R.S., Martinou, J.C., 2012. Identification and functional expression of the mitochondrial pyruvate carrier. Science (1979) 336, 93–96. 10.1126/science.1218530

Janse, C.J., Ramesar, J., Waters, A.P., 2006. High-efficiency transfection and drug selection of genetically transformed blood stages of the rodent malaria parasite Plasmodium berghei. Nat Protoc 1, 346–356. 10.1038/nprot.2006.53

Jeffers, V., Child, M.A., 2022. Previews No acetyl-CoA keeps Plasmodium at bay. Cell Chem Biol 29, 174–176. 10.1016/j.chembiol.2022.02.003

Jones, D.T., Taylor, W.R., Thornton, J.M., 1992. The rapid generation of mutation data matrices from protein sequences. Bioinformatics 8, 275–282. 10.1093/bioinformatics/8.3.275

Jortzik, E., Fritz-Wolf, K., Sturm, N., Hipp, M., Rahlfs, S., Becker, K., 2010. Redox regulation of plasmodium falciparum ornithine δ-aminotransferase. J Mol Biol 402, 445–459. 10.1016/j.jmb.2010.07.039

Ke, H., Lewis, I.A., Morrisey, J.M., McLean, K.J., Ganesan, S.M., Painter, H.J., Mather, M.W., Jacobs-Lorena, M., Llinás, M., Vaidya, A.B., 2015. Genetic investigation of tricarboxylic acid metabolism during the plasmodium falciparum life cycle. Cell Rep 11, 164–174. 10.1016/j.celrep.2015.03.011

Koper, K., Hataya, S., Hall, A. G., Takasuka, T. E., & Maeda, H. A. (2023). Chapter Two - Biochemical characterization of plant aromatic aminotransferases. In J. Jez (Ed.), Methods in Enzymology (Vol. 680, pp. 35–83). Academic Press. 10.1016/bs.mie.2022.07.034

Kronenberger, T., Lindner, J., Meissner, K.A., Zimbres, F.M., Coronado, M.A., Sauer, F.M., Schettert, I., Wrenger, C., 2014. Vitamin B6-dependent enzymes in the human malaria parasite plasmodium falciparum: A druggable target? Biomed Res Int. 108516. 10.1155/2014/108516.

Li, Xiaoli, Ji, Y., Han, G., Li, Xiaoran, Fan, Z., Li, Y., Zhong, Y., Cao, J., Zhao, J., Zhang, M., Wen, J., Goscinski, M.A., Nesland, J.M., Suo, Z., 2016. MPC1 and MPC2 expressions are associated with favorable clinical outcomes in prostate cancer. BMC Cancer 1–12. 10.1186/s12885-016-2941-6

Le, X.H., Lee, C.P., Harvey Millar, A., 2021. The mitochondrial pyruvate carrier (MPC) complex mediates one of three pyruvate-supplying pathways that sustain Arabidopsis respiratory metabolism. Plant Cell 33, 2776–2793. 10.1093/plcell/koab148

Lyu, C., Chen, Y., Meng, Y., Yang, J., Ye, S., Niu, Z., EI-Debs, I., Gupta, N., Shen, B., 2023. The Mitochondrial Pyruvate Carrier Coupling Glycolysis and the Tricarboxylic Acid Cycle Is Required for the Asexual Reproduction of Toxoplasma gondii. Microbiol Spectr. 11. 10.1128/spectrum.05043-22

MacKay, G.M., Zheng, L., Van Den Broek, N.J.F., Gottlieb, E., 2015. Analysis of Cell Metabolism Using LC-MS and Isotope Tracers. Methods Enzymol 561, 171–196. 10.1016/bs.mie.2015.05.016

MacRae, J.I., Dixon, M.W.A., Dearnley, M.K., Chua, H.H., Chambers, J.M., Kenny, S., Bottova, I., Tilley, L., McConville, M.J., 2013. Mitochondrial metabolism of sexual and asexual blood stages of the malaria parasite Plasmodium falciparum. BMC Biol 11. 10.1186/1741-7007-11-67

MacRae, J.I., Sheiner, L., Nahid, A., Tonkin, C., Striepen, B., McConville, M.J., 2012. Mitochondrial metabolism of glucose and glutamine is required for intracellular growth of toxoplasma gondii. Cell Host Microbe 12, 682–692. 10.1016/j.chom.2012.09.013

Manzoni, G., Briquet, S., Risco-Castillo, V., Gaultier, C., Topçu, S., Ivǎnescu, M.L., Franetich, J.F., Hoareau-Coudert, B., Mazier, D., Silvie, O., 2014. A rapid and robust selection procedure for generating drug-selectable marker-free recombinant malaria parasites. Sci Rep 4, 1–10. 10.1038/srep04760

Martínez-Reyes, I., Chandel, N.S., 2020. Mitochondrial TCA cycle metabolites control physiology and disease. Nat Commun 11, 1–11. 10.1038/s41467-019-13668-3

Nagappa, L.K., Singh, D., Dey, S., Kumar, K.A., Balaram, H., 2019. Biochemical and physiological investigations on adenosine 5ʹ monophosphate deaminase from Plasmodium spp. Mol Microbiol. 112, 699–717. 10.1111/mmi.14313

Nair, S.C., Munro, J.T., Mann, A., Llinás, M., Prigge, S.T., 2023. The mitochondrion of Plasmodium falciparum is required for cellular acetyl-CoA metabolism and protein acetylation. Proc Natl Acad Sci U S A 120, e2210929120. 10.1073/PNAS.2210929120

Negreiros, R.S., Lander, N., Chiurillo, M.A., Vercesi, A.E., Docampo, R., 2021. Mitochondrial pyruvate carrier subunits are essential for pyruvate-driven respiration, infectivity, and intracellular replication of trypanosoma cruzi. mBio 12. 10.1128/MBIO.00540-21.

Oppenheim, R.D., Creek, D.J., Macrae, J.I., Modrzynska, K.K., Pino, P., Limenitakis, J., Polonais, V., Seeber, F., Barrett, M.P., Billker, O., Mcconville, M.J., Soldati-favre, D., 2014. BCKDH: The Missing Link in Apicomplexan Mitochondrial Metabolism Is Required for Full Virulence of Toxoplasma gondii and Plasmodium berghei 10. 10.1371/journal.ppat.1004263

Orr, R.Y., Philip, N., Waters, A.P., 2012. Improved negative selection protocol for Plasmodium berghei in the rodent malarial model. Malar J 11, 8–13. 10.1186/1475-2875-11-103

Papa, S., Francavilla, A., Paradies, G., Meduri, B., 1971. The transport of pyruvate in rat liver mitochondria. FEBS Lett 12, 285–288. 10.1016/0014-5793(71)80200-4

Patil, H., Hughes, K.R., Lemgruber, L., Philip, N., Dickens, N., Lucas Starnes, G., Waters, A.P., 2020. Zygote morphogenesis but not the establishment of cell polarity in Plasmodium berghei is controlled by the small GTPase, RAB11A. PLoS Pathog 16, 1–27. 10.1371/journal.ppat.1008091

Pfander, C., Anar, B., Schwach, F., Otto, T.D., Brochet, M., Volkmann, K., Quail, M.A., Pain, A., Rosen, B., Skarnes, W., Rayner, J.C., Billker, O., 2011. A scalable pipeline for highly effective genetic modification of a malaria parasite. Nat Methods 8, 1078–1084. 10.1038/nmeth.1742

Pietrocola, F., Galluzzi, L., Bravo-San Pedro, J.M., Madeo, F., Kroemer, G., 2015. Acetyl coenzyme A: a central metabolite and second messenger. Cell Metab 21, 805–821. 10.1016/J.CMET.2015.05.014

Prata, I.O., Cubillos, E.F.G., Krüger, A., Barbosa, D., Martins, J., Setubal, J.C., Wunderlich, G., 2021. Plasmodium falciparum Acetyl-CoA Synthetase Is Essential for Parasite Intraerythrocytic Development and Chromatin Modification. ACS Infect Dis 7, 3224– 3240. 10.1021/acsinfecdis.1c00414

Quang, L.S., Gascuel, O., Lartillot, N., 2008. Empirical profile mixture models for phylogenetic reconstruction. Bioinformatics 24, 2317–2323. 10.1093/bioinformatics/btn445

Rajaram, K., Rangel, G.W., Munro, J.T., Nair, S.C., Llinás, M., Prigge, S.T., 2024. MULTIPLE, REDUNDANT CARBOXYLIC ACID TRANSPORTERS SUPPORT MITOCHONDRIAL METABOLISM IN PLASMODIUM FALCIPARUM. bioRxiv 2024.11.26.624872. 10.1101/2024.11.26.624872

Rodríguez, M.C., Margos, G., Compton, H., Ku, M., Lanz, H., Rodríguez, M.H., Sinden, R.E., 2002. Plasmodium berghei: Routine production of pure gametocytes, extracellular gametes, zygotes, and ookinetes. Exp Parasitol 101, 73–76. 10.1016/S0014-4894(02)00035-8

Schindelin, J., Arganda-Carrera, I., Frise, E., Verena, K., Mark, L., Tobias, P., Stephan, P., Curtis, R., Stephan, S., Benjamin, S., Jean-Yves, T., Daniel, J.W., Volker, H., Kevin, E., Pavel, T., Albert, C., 2009. Fiji - an Open platform for biological image analysis. Nat Methods 9. 10.1038/nmeth.2019.Fiji

Schneider, C.A., Rasband, W.S., Eliceiri, K.W., 2012. NIH Image to ImageJ: 25 years of image analysis. Nat Methods 9, 671–675. 10.1038/nmeth.2089

Schwach, F., Bushell, E., Gomes, A.R., Anar, B., Girling, G., Herd, C., Rayner, J.C., Billker, O., 2015. PlasmoGEM, a database supporting a community resource for large-scale experimental genetics in malaria parasites. Nucleic Acids Res 43, D1176–D1182. 10.1093/nar/gku1143

Seeber, F., Limenitakis, J., Soldati-Favre, D., 2008. Apicomplexan mitochondrial metabolism: a story of gains, losses and retentions. Trends Parasitol. 24, 468–478. 10.1016/j.pt.2008.07.004

Shi, L., Tu, B.P., 2015. Acetyl-CoA and the regulation of metabolism: Mechanisms and consequences. Curr Opin Cell Biol 33, 125–131. 10.1016/j.ceb.2015.02.003

Srivastava, A., Creek, D.J., Evans, K.J., De Souza, D., Schofield, L., Müller, S., Barrett, M.P., McConville, M.J., Waters, A.P., 2015. Host Reticulocytes Provide Metabolic Reservoirs That Can Be Exploited by Malaria Parasites. PLoS Pathog 11, 1–22. 10.1371/journal.ppat.1004882

Srivastava, A., Philip, N., Hughes, K.R., Georgiou, K., MacRae, J.I., Barrett, M.P., Creek, D.J., McConville, M.J., Waters, A.P., 2016. Stage-Specific Changes in Plasmodium Metabolism Required for Differentiation and Adaptation to Different Host and Vector Environments. PLoS Pathog 12, 1–30. 10.1371/journal.ppat.1006094

Štáfková, J., Mach, J., Biran, M., Verner, Z., Bringaud, F., Tachezy, J., 2016. Mitochondrial pyruvate carrier in Trypanosoma brucei. Mol Microbiol 100, 442–456. 10.1111/MMI.13325

Summers, R.L., Pasaje, C.F.A., Pisco, J.P., Striepen, J., Luth, M.R., Kumpornsin, K., Carpenter, E.F., Munro, J.T., Lin, D., Plater, A., Punekar, A.S., Shepherd, A.M., Shepherd, S.M., Vanaerschot, M., Murithi, J.M., Rubiano, K., Akidil, A., Ottilie, S., Mittal, N., Dilmore, A.H., Won, M., Mandt, R.E.K., McGowen, K., Owen, E., Walpole, C., Llinás, M., Lee, M.C.S., Winzeler, E.A., Fidock, D.A., Gilbert, I.H., Wirth, D.F., Niles, J.C., Baragaña, B., Lukens, A.K., 2022. Chemogenomics identifies acetyl-coenzyme A synthetase as a target for malaria treatment and prevention. Cell Chem Biol 29, 191–201.e8. 10.1016/j.chembiol.2021.07.010

Tamura, K., Stecher, G., Kumar, S., 2021. MEGA11: Molecular Evolutionary Genetics Analysis Version 11. Mol Biol Evol 38, 3022–3027. 10.1093/molbev/msab120

Tavoulari, S., Schirris, T.J.J., Mavridou, V., Thangaratnarajah, C., King, M.S., Jones, D.T.D., Ding, S., Fearnley, I.M., Kunji, E.R.S., 2022. Key features of inhibitor binding to the human mitochondrial pyruvate carrier hetero-dimer. Mol Metab 60, 101469. 10.1016/j.molmet.2022.101469

Tavoulari, S., Thangaratnarajah, C., Mavridou, V., Harbour, M.E., Martinou, J., Kunji, E.R., 2019. The yeast mitochondrial pyruvate carrier is a hetero-dimer in its functional state. EMBO J 38, 1–13. 10.15252/embj.2018100785

Venkatesan, M., Amaratunga, C., Campino, S., Auburn, S., Koch, O., Lim, P., Uk, S., Socheat, D., Kwiatkowski, D.P., Fairhurst, R.M., Plowe, C. V., 2012. Using CF11 cellulose columns to inexpensively and effectively remove human DNA from Plasmodium falciparum-infected whole blood samples. Malar J 11, 1–7. 10.1186/1475-2875-11-41

Voet, D., Voet, J.G. and Pratt, C.W., 2016. Fundamentals of biochemistry: Life at the molecular level. 5^th^ edition. WILEY publications.

Wrenger, C., Müller, I.B., Schifferdecker, A.J., Jain, R., Jordanova, R., Groves, M.R., 2011. Specific inhibition of the aspartate aminotransferase of plasmodium falciparum. J Mol Biol 405, 956–971. 10.1016/j.jmb.2010.11.018

Xu, Y., Tao, Y., Cheung, L.S., Fan, C., Chen, L.Q., Xu, S., Perry, K., Frommer, W.B., Feng, L., 2014. Structures of bacterial homologues of SWEET transporters in two distinct conformations. Nature 515, 448–452. 10.1038/nature13670

Zangari, J., Petrelli, F., Maillot, B., Martinou, J.C., 2020. The multifaceted pyruvate metabolism: Role of the mitochondrial pyruvate carrier. Biomolecules 10, 1–18. 10.3390/biom10071068

