## Supplementary material for "Understanding the biochemical significance of the mitochondrial pyruvate carrier in the rodent malarial parasite, *Plasmodium berghei*": Supple_MPC_bioRxiv: Supple_MPC_bioRxiv.pdf

**Composition of the gametocyte medium.**
For 100 mL of gametocyte medium, 1.04 g of RPMI-1640 (10.4 g /L), 594 mg of HEPES (25 mM), 200 mg of $NaHCO_3$ (23.8 mM), and 1 mL of 100X HT (Gibco, mixture of 10mM sodium hypoxanthine and 1.6 mM thymidine) supplement was dissolved in 80 mL of sterile water. The pH of the solution was adjusted to 7.3 and the volume was made up to 100 mL. The solution was filtered using a 0.2 µm sterile filter and stored at 4 ℃ after the addition of 100 µL of 40 mg /L gentamycin.

**Composition of the minimal medium**
The composition of minimal medium was adapted from Srivastava A (2014, http://theses.gla.ac.uk/5725/)
For every 100 mL of minimal medium, the following components were added sequentially.
1) 0.5 g Alubmax™ I (5 g /L) and 6.5 mg of L-cystine.2HCl were added to 80 mL of solution A. 2) 10 mL of solution B (amino acids, 10x stock). 3) 10 µL of solution C (calcium pantothenate, 1000x stock). 4) 1 mL of solution D (hypoxanthine, 100x stock). 5) 1 mL of solution E (glutathione, 100x stock). 6) 5 mL of solution F (D-glucose, 20x stock). 7) 1 mL of solution G (phenol red, 100x stock).
The volume was made up to 100 mL by adding 2 mL of sterile water and pH was adjusted to 7.3. The media was filter sterilized using a 0.22 µm membrane filter and 100 µL of gentamycin (40 mg /mL) was added under sterile conditions and stored at 4 ℃.
A 100 mL of minimal medium without D-glucose was also prepared simultaneously and filtered using a 0.2µm sterile filter. 100 µL of gentamycin (40 mg /mL) was added and kept at 4 ℃ before use.

*Solution A (Salts and other components [1x stock]).* Given in Table A1 is the list of components that were weighed and dissolved sequentially in 400 mL of sterile water. The solution was filter-sterilized using a 0.2 µm membrane filter. Aliquots of 50 mL in sterile falcon tubes were stored at -20 ℃. It should be noted that the volume was intentionally not adjusted to 500 mL. The components that were dissolved in 400 mL had a stock concentration of 1.25x. They were adjusted to a final concentration of 1x when used for 100 mL of minimal media preparation. Hence, 80 mL of this salt solution (1.25x stock) was used for every 100 mL of minimal medium.

Solution A: salts and buffer (1.25x stock)

| **Components** | **mg/ 400 mL** |
|---|---|
| KCl | 200 mg |
| NaCl | 2650 mg |
| $Na_2HPO_4$ (anhydrous) | 400 mg |
| $NaHCO_3$ | 1000 mg |
| $MgSO_4$ (anhydrous) | 24.4 mg |
| $Ca(NO_3)_2.4H20$ | 50 mg |
| HEPES | 2979 mg |

*Solution B: Amino acids [10x stock].* 250 mL of a 10x stock solution of amino acids was prepared as described earlier (Srivastava A, 2014, http://theses.gla.ac.uk/5725/). The solution was filter-sterilized and stored at 4 ℃. Details of stock solution preparation that need attention are enumerated below.

- A 10x stock solution of L-cystine.2HCl (Sigma-Aldrich) was not prepared due to its poor solubility. This amino acid was directly added to a final concentration of 6.5 mg / 100 mL into the complete minimal medium.

- For culturing with $^{13}C$ L-glutamine, 100 mL of a set of amino acids (Table below) at 10x concentration without $^{12}C$ L-glutamine was made, filter-sterilized, and stored at 4 ℃.

- For labelling experiments with glutamine, 5 mL of 146.14 mM U-$^{13}C_5$ $^{15}N_2$ glutamine (20x stock) solution was made. For this, 29.22 mg of U-$^{13}C_5$ $^{15}N_2$ glutamine was dissolved in 5 mL of sterile water, filter-sterilized, and aliquots of 1 mL were stored at -80 ℃.

**Solution B: 10x stock of amino acids**

| Components | mg / 250 mL |
|---|---|
| L-glutamic acid | 50 mg |
| L-glutamine | 750 mg |
| L-Isoleucine | 125 mg |
| L-methionine | 37.5 mg |
| L-Proline | 50 mg |
| L-Tyrosine disodium dihydrate | 75 mg |

*Solution C: Calcium pantothenate [10,000x stock].* To prepare 10mL of 10,000x stock solution, 25 mg of calcium pantothenate (Sigma Aldrich) was dissolved in 10 mL of sterile water, filter sterilized using 0.22 µm membrane filter, and stored at 4 ℃.

*Solution D: Hypoxanthine [100x stock].* For 50 mL of 100x stock solution, 20.5 mg of hypoxanthine (Sigma Aldrich) was dissolved in 50 mL of sterile water containing 0.35N NaOH. The solution was filter sterilized, and aliquots of 1 mL were stored at -20 ℃.

*Solution E: Glutathione [100x stock].* For 50 mL of 100x stock solution, 5 mg of reduced L-glutathione (Sigma Aldrich) was dissolved in 50 mL of sterile water, filter-sterilized, and aliquots of 1 mL were stored at 4 ℃.

*Solution F: D-Glucose [20x stock].* 50 mL of 160 mM D-glucose (20x stock) solution was prepared by dissolving 1.4 g of D-glucose (Sigma Aldrich) in 50 mL sterile water. The solution was filter-sterilized and stored at 4 ℃.

For labelling experiments with glucose, 148.88 mg of $^{13}C_6$-D-glucose was dissolved in 5 mL of sterile water, filter-sterilized and 1 mL aliquots were stored at -80 ℃.

*Solution G: Phenol red [100x stock].* For 10 mL of 100x stock solution, 5 mg of Phenol Red dye (Sigma Aldrich), a water-soluble pH indicator was dissolved in 10 mL of sterile water, filter-sterilized, and stored at 4 ℃.

**Table 1. Primers used for PCRs.**

| S.N. | Primer name | Primer sequence (5' to 3') |
|---|---|---|
| **Primers used for the generation of knockout vectors and genotyping of Δ*mpc1*, Δ*mpc2*, and Δ*mpc1mpc2* parasites** | | |
| P1 | PbMPC1-QCR1 FP | GGAAAAGAATGTATTACGAATTGAAACCTTC |
| P2 | PbMPC1-QCR2 RP | CAATAACAAAACCCCAATTTGCTAAGG |
| P3 | PbMPC1-5'int FP | CTGCCAGCGGGATAGTTCTTGTAG |
| P4 | PbMPC1-3'int RP | CGTGTTAAACCGCGTGCTTCATGTGATATGGC |
| P5 | p413PbMPC1_ FP | GCCGGAATTCATGTCAAAATTTAAATTACTTTTTCAAAATGTAAAG |
| P6 | p413PbMPC1_ RP | GCGGGTCGACTTAGTGATGGTGATGGTGATGACCAGATCCACCGTTGGTTG CTAACATCAATTTATTATCTG |
| P7 | PbMPC2 QCR1 FP | CATGGTAAATATGATATGAAAGAATAAG |
| P8 | PbMPC 2QCR2 RP | GTGTGCGCATATATGGAGCT TAAATG |
| P9 | PbMPC2- 5' int FP | GACGACAATCAAGGGCTGCAAAATAGTAATC |
| P10 | PbMPC2-3' int RP | GAGCCAAATAGGAATGGTAAAGTGAATGGG |
| P11 | p416 PbMPC2 FP | GCCGGGATCCATGAATATAATTAGAAAAGTTTTTTATCCAAATATTG |
| P12 | p416 PbMPC2 RP | GCGGGTCGACTTAGTGATGGTGATGGTGATGACCAGATCCACCCTCTTTAA TTTCCTTCTCGTTGTTTATG |
| P13 | hDHFR-Yfcu | ACTTCTTAAACCTAATCTGTAGTAAGGAAGGGATTG |
| P14 | Pbef1alpha promoter FP | GAATATTAAATTGTAAACTTAAGCATAAAGAGCTCG |
| P15 | Yfcu FP | CATCTAGACCTCTGTCGAGGGCACCAGTAACAATTCTG |
| P16 | hDHFR RP | CCACAACCTCTTCAGTAGAAGGTAAACAGAATCTGGTG |
| P17 | PbMPC2 3'flank RP | GGTTCATATTTATTTAGTACTTTACGC |
| **Primers used for the generation of PbMPC1GFPDDD1XHA and PbMPC2GFPDDD1XHA tagging constructs** | | |
| P18 | MPC2 tag-rec2 RP | AATTAAAAAAAAACATTTATTTATTCCTTCTTTTATAATTTATTGTCCTTTCCG CCTACTGCGACTATAGA |
| P19 | MPC2 tag-rec1 FP | TTGCCGTATATAAATATAATAACATAAACAACGAGAAGGAAATTAAAGAG AAGGCGCATAACGATACCAC |
| P20 | MPC2-FP | AATTAAAAAAAAACATTTATTTATTCCTTCTTTTATAATTTATTGTCCTTTAA GGCGCATAACGATACCAC |
| P21 | MPC2-3'flank (RP) | GGTTCATATTTATTTAGTACTTTACGC |
| P22 | MPC1 tag-rec R1 FP | TTCGAAAGTTAAAATACGGTTCAGATAATAAATTGATGTTAGCAACCAACA AGGCGCATAACGATACCAC |
| P23 | MPC1 tag-rec R2 RP | AGGTGATTAATTTTTAGGTATGTCCTTTGTTTACATGATTAGGTATGTTTCC GCCTACTGCGACTATAGA |
| P24 | MPC1-3'flank RP | CGCAAGTTTATAAAAAATTGGGTAC |
| P25 | MPC1 FP | GAGAAAATGACATCAGTCTTAGTTG |
| P26 | Zeo/PheS RP | TCATTCTTCGAAAACGATCTGCG |
| P27 | GFP internal RP | CAATGCTTTGCGAGATACCCAGATCATATG |
| P28 | Yfcu_1 FP | GGCAAGCAAGTGGGATCAGAAGGGTATGGACATTGC |
| P29 | HA tag RP | AGCGTAATCTGGAACATCGTATGGGTA |
| **Primers used for the validation of yeast *Scmpc1*Δ, *Scmpc2*Δ, *Scmpc2*Δ*mpc3*Δ and *Scmpc1*Δ*mpc2*Δ*3*Δ strains** | | |
| P30 | 5'UTR scMPC1 FP | GCTTTAACCGAGGCGAAATGACAAG |
| P31 | 3'UTR scMPC1 RP | GTCATCTTATGTAGTTTGGTTCAATAC |
| P32 | 5'UTR scMPC2 FP | GTTGACATCGCTGACTGCAATAG |
| P33 | 3'UTR scMPC2 RP | GCAAAAGAGCAAATTATAACACCG |
| P34 | 5'UTR scMPC3 FP | GATAATATGGTGCGTTGCAATTGG |
| P35 | 3'UTR scMPC3 RP | GACCCAAAATTAAGCTTCTCGAG |
| **Primers used for generation of C-terminal 6x-His tagged constructs in yeast expression plasmids** | | |
| P5 | p413PbMPC1_EcoRI FP | GCCGGAATTCATGTCAAAATTTAAATTACTTTTTCAAAATGTAAAG |
| P6 | p413PbMPC1_EcoRI RP | GCGGGTCGACTTAGTGATGGTGATGGTGATGACCAGATCCACCGTTGGTTG CTAACATCAATTTATTATCTG |
| P11 | p416PbMPC2_BamH IFP | GCCGGGATCCATGAATATAATTAGAAAAGTTTTTTATCCAAATATTG 3' |
| P12 | p416PbMPC2_SalI RP | GCGGGTCGACTTAGTGATGGTGATGGTGATGACCAGATCCACCCTCTTTAA TTTCCTTCTCGTTGTTTATG |
| P36 | p413ScMPC1_EcoRI FP | GCCGGAATTCATGTCTCAACCGGTTCAACGCGCTGC |
| P37 | p413ScMPC1_SalI RP | GCGGGTCGACTTAGTGATGGTGATGGTGATGACCAGATCCACCCTGTTTAC CAGTTTTTTCTTTCTCTTTCCATTCC |
| P38 | p416ScMPC2_SalI RP | GCTGGTCGACTTAGTGATGGTGATGGTGATGACCAGATCCACCTCTGCCCG TAGTAATTTCCTTTTTGCTTTC |

| | | |
|---|---|---|
| P39 | p416ScMPC2_BamHI FP | GCGGGGATCCATGTCTACATCATCCGTACGTTTTGCATTTAGG |
| **Primer set used as internal positive control in genotyping of wild-type, Δ*mpc1*, Δ*mpc2*, and Δ*mpc1mpc2* parasites** | | |
| P53 | PbDTC FP | GGATATGCCCCAAATTAATTATGCATGC |
| P54 | PbDTC RP | GGTACGAAATAAACCTAATCGTCCTGTTG |

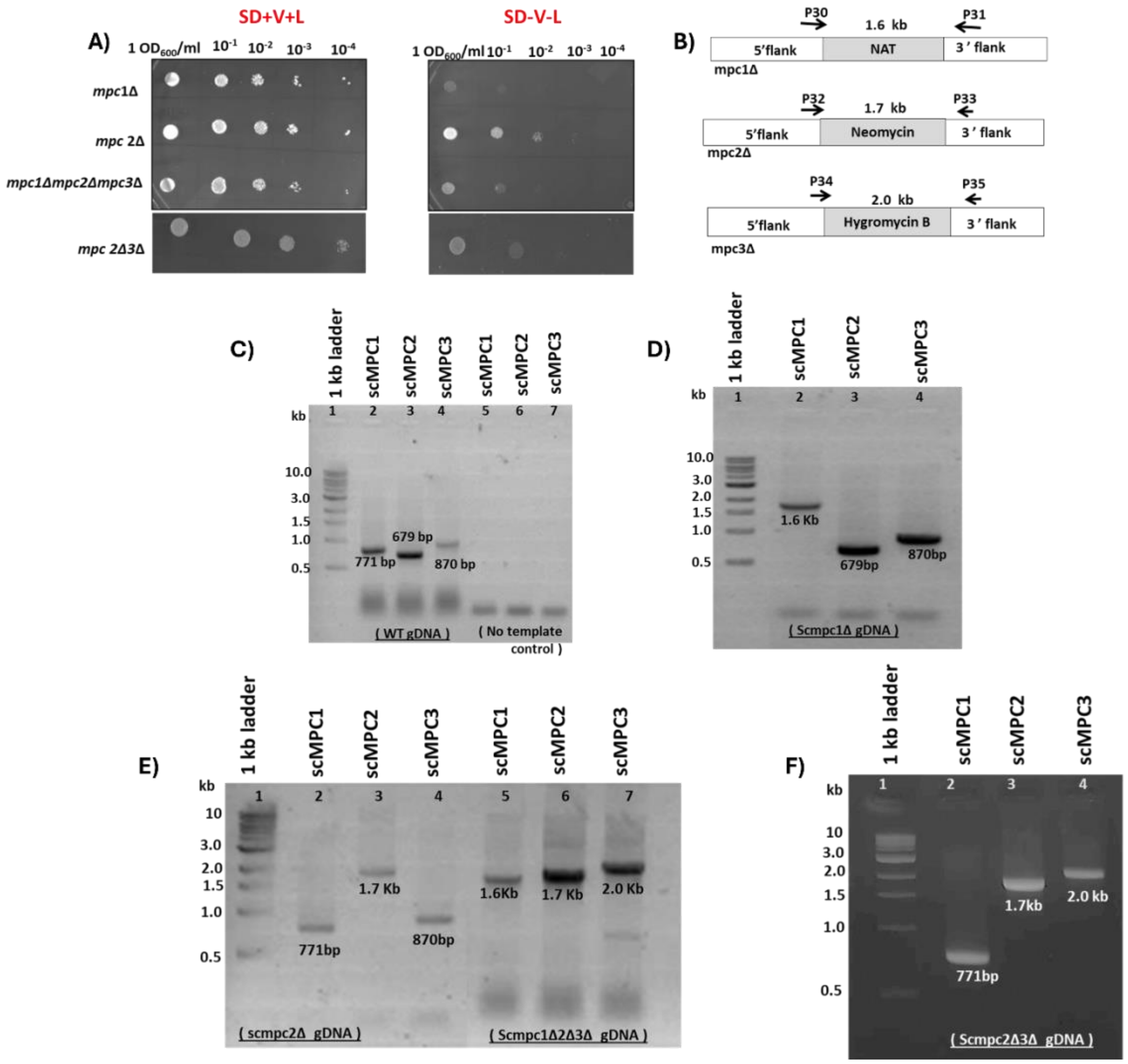

**Figure S1. Validation of ScΔmpc1, ScΔmpc2, ScΔmpc2Δmpc3 and ScΔmpc1Δmpc2Δmpc3 strains of S. cerevisiae.** A) Spot assay for the confirmation of *Δmpc1, Δmpc2, Δmpc2Δmpc3, and Δmpc1Δmpc2Δmpc3* yeast strains. The absence of functional MPC will inhibit the growth of *mpc* deficient yeast strains in minimal medium lacking leucine and valine. B) A schematic illustration of *Δmpc1, Δmpc2, and Δmpc3* gene locus. The indicated primers were used to confirm *mpc* deleted strains by PCR. Genomic DNA was isolated from wild-type or *Δmpc* yeast cells and used as the template for PCR genotyping. C) PCR was carried out to confirm the presence of Sc*mpc1*, Sc*mpc2*, and Sc*mpc3* genes using wild-type gDNA as template. This served as our positive control whereas PCR without template served as a negative control. PCR validation of Sc*Δmpc1*, Sc*Δmpc2* and Sc*Δmpc2Δmpc3*, and Sc*Δmpc1Δmpc2Δmpc3* strains are shown in (D), (E), and (F), respectively. The deletion of the Sc*mpc1*, Sc*mpc2*, or Sc*mpc3* genes was confirmed by using primer pairs indicated in panel B. The amplified DNA fragments of the expected size, 1.6 kbp (Sc*Δmpc1)*, 1.7 kbp (Sc*Δmpc2)*, and 2 kbp (ScΔmpc3) were observed for the different knockout lines.

A)

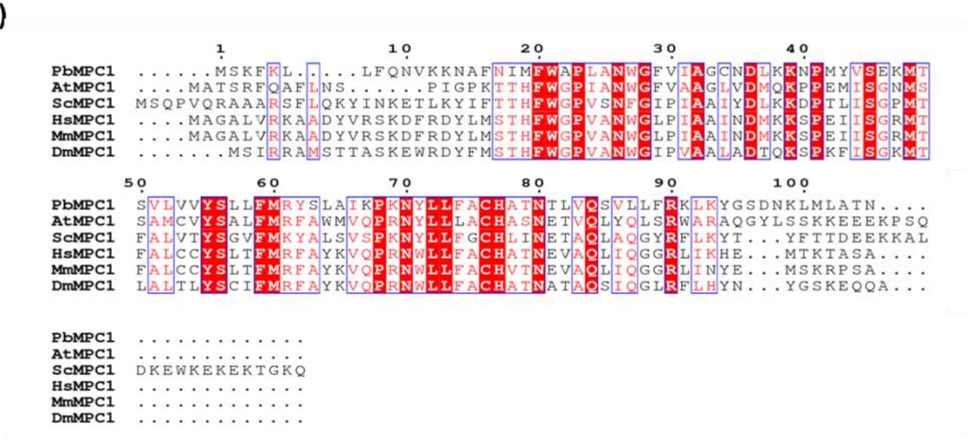

B)

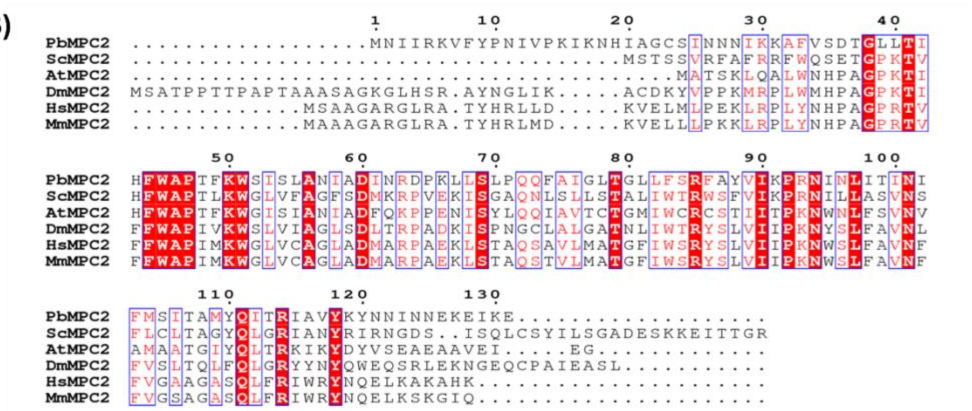

**Figure S2. Multiple sequence alignment of P. berghei MPC1 (A) and MPC2 (B)** protein sequences with orthologs from *Arabidopsis thaliana*, *Saccharomyces cerevisiae*, *Homo sapiens*, *Mus musculus* and *Drosophila melanogaster*. Alignment was generated using Clustal Omega and rendered using ESpript 3.0. The red-shaded segments highlight residues invariant across the different organisms.

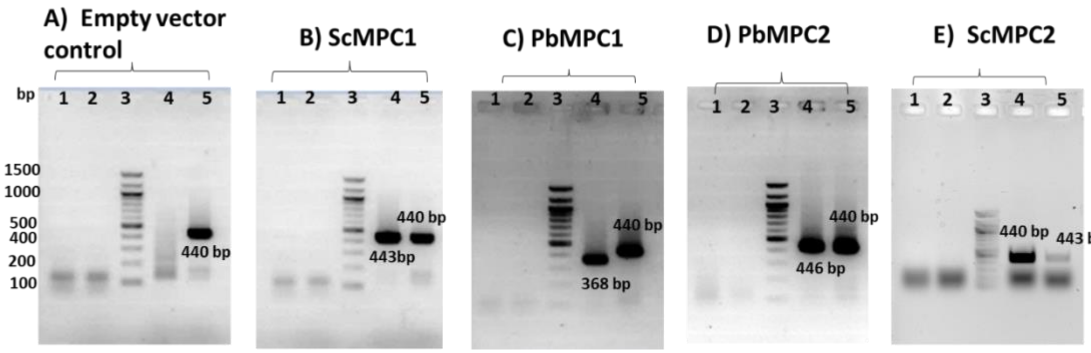

**Figure S3. Validation of the expression of Sc*mpc1*, Sc*mpc2*, Pb*mpc1*, and Pb*mpc2* genes in *S. cerevisiae* at the transcript level by RT-PCR**. Agarose gel electrophoresis of RT-PCR mixtures confirms transcription of Sc*mpc1*, Pb*mpc1*, and Pb*mpc2* in transformed *S. cerevisiae* ***Δ***mpc1 cells carrying the plasmid p413_ScMPC1 (panel B), p413_PbMPC1 (panel C), and p416_PbMPC2 (panel D) plasmids, respectively. The transcription of ScMPC2 was confirmed in *S. cerevisiae Δmpc2Δmpc3* cells carrying the plasmid p416-ScMPC2 (panel E). In all panels (A-E), lanes 1 and 2 correspond to controls wherein DNase I treated RNA isolated from plasmid transformed *S. cerevisiae Δmpc1* or *Δmpc2Δmpc3* cells was not subjected to reverse transcription reaction. This control confirms the absence of genomic DNA contamination. Lane 3, 100 bp DNA ladder. Lane 4, RT-PCR with RNA from the indicated gene transformed into ***Δ***mpc1yeast cells. Lane 5, RT-PCR of ScMPC2 (440 bp) RNA from *Δmpc1* and of ScMPC1 (443 bp) RNA from *Δmpc2Δmpc3* served as internal positive controls. RT-PCR with RNA from Δmpc1 cells carrying empty vectors served as a negative control (panel A).

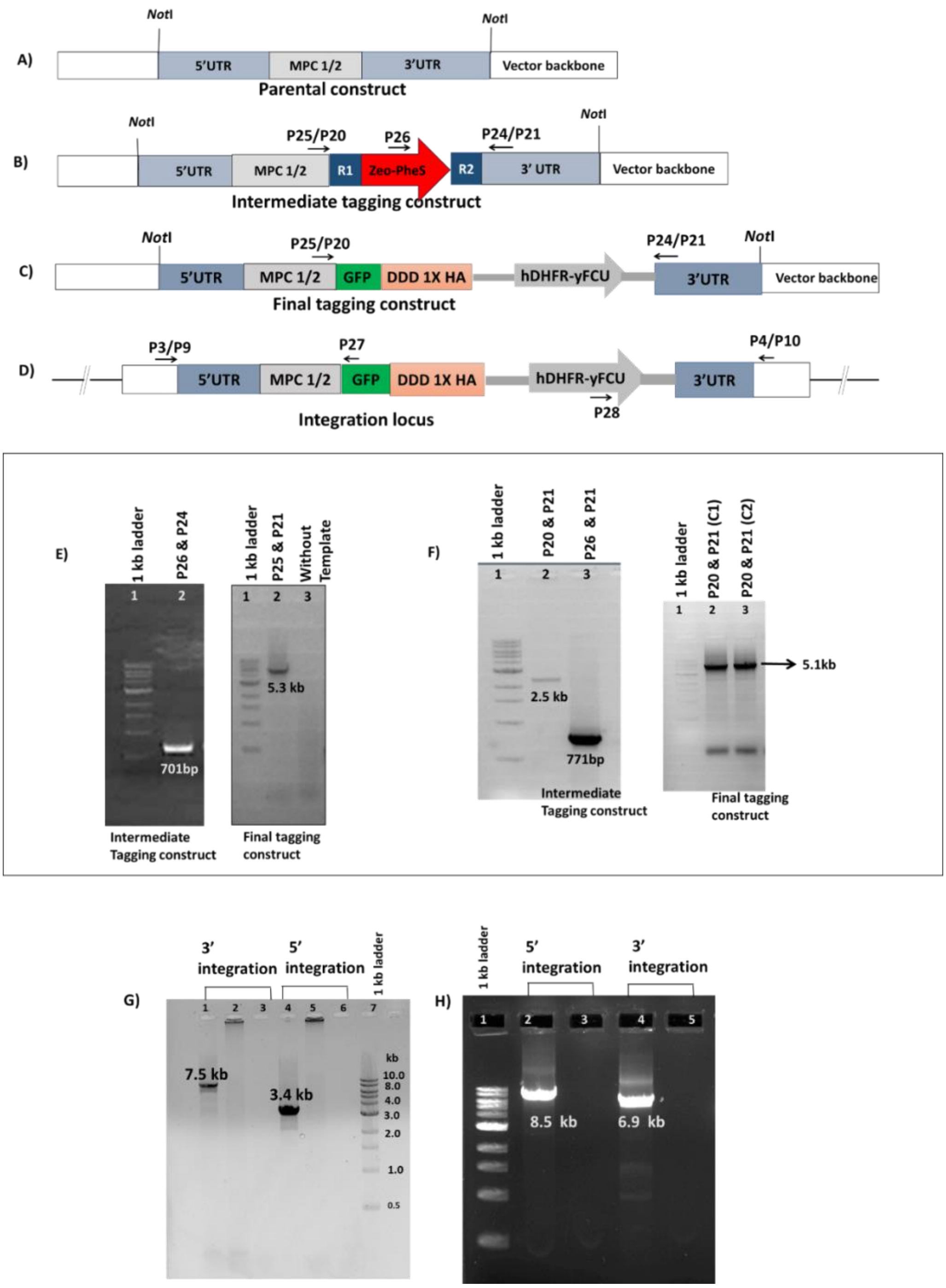

**Figure S4. Generation of transgenic parasite lines expressing GFP-DDD-HA tagged PbMPC1 and PbMPC2 proteins.** (A-C) Schematic of the parental, intermediate, and tagging constructs of PbMPC1_GFP-DDD-1xHA and PbMPC2_GFP-DDD-1xHA (DDD, DHFR destabilization domain). D) Locus of the tagged *mpc1* or *mpc2* genes in the parasite genome after integration through homologous double crossover recombination. E) Agarose gel electrophoresis of PCR reactions carried out to confirm PbMPC1 intermediate tagging constructs. Left panel, confirmation of PbMPC1 intermediate tagging construct by PCR using the indicated primers. Lane 1, 1 kbp ladder; Lane 2, PCR validation using P26 and P24 as primers. Right panel, confirmation of PbMPC1 final tagging construct. Lane 1, 1 kb ladder; PCR validation using P25 and P21 primers with template (lane 2) and without template (lane 3). F) Agarose gel electrophoresis of PCR reactions carried out to confirm PbMPC2 intermediate and tagging construct. Left panel, validation of the intermediate tagging construct. Lane 1, 1 kb ladder; Lane 2 PCR confirmation using primer pairs; P20 & P21, and P26 & P21. The right panel confirms the generation of the

PbMPC2 final tagging construct. Two clones; C1 and C2 were confirmed. Lane 1, 1 kb ladder; Lanes 2-3, PCRs using P20 and P21 as primers. (G-H) Genotyping of PbMPC1-GFP-DDD-1XHA (G) and PbMPC2-GFP-DDD-1XHA (H) tagged parasites. Primers P3 and P4 for MPC1, P9 and P10 for MPC2, P27 and P28 specific for GFP and hDHFR used for PCR genotyping are indicated. G) PCR with primers P4 and 28 to confirm 3' integration using genomic DNA from GFP-tagged PbMPC1 parasites (lane 1), WT (lane 2), and without template (lane 3). PCR with primers P3 and P27, to confirm 5' integration using gDNA from GFP tagged PbMPC1 parasites (lane 4), WT (lane 5), and without template (lane 6). H) PCR with primers P9 and P27 to confirm 5' integration using gDNA from GFP-tagged PbMPC2 parasites (lane 1) and without template (lane 2). PCR with primers P10 and P28, to confirm 3' integration using gDNA from GFP tagged PbMPC2 parasites (lane 3) and without template (lane 4). All amplified products of PCR showed bands of the expected size. kb in all panels should be read as kbp (kilo base pairs).

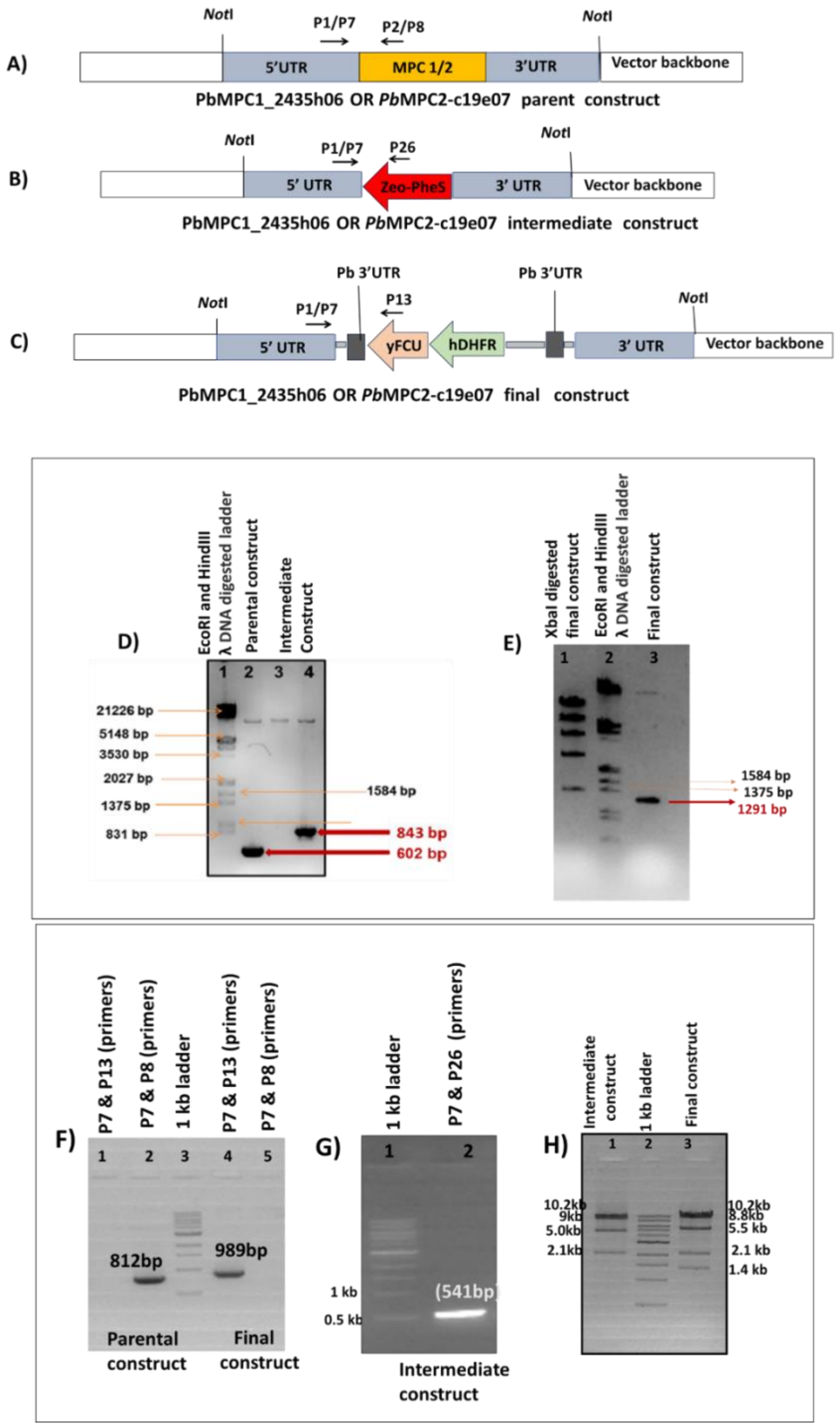

**Figure S5. Generation of PbMPC1 and PbMPC2 final knockout construct using recombineering-based methodology. (A-C)** Schematic of pJAZZPbMPC1_2435h06 or pJAZZPbMPC2_c19e07 parental, intermediate, and final knockout constructs. The oligonucleotides used for the confirmation of these constructs and their positions are indicated as arrows. (D-E) PCR Confirmation of pJAZZPbMPC1_2435h06 parental, intermediate, and final knockout constructs. D) Lane 1, HindIII + EcoRI digested λ DNA marker; lane 2, PCR validation of the parental construct using P1 and P2 as primers. Confirmation of the intermediate construct by PCR using P1 and P26 as primers: without template (lane 3) and with template (lane 4). E) Lane1, XbaI restriction digestion of final knockout construct; Lane 2, HindIII + EcoRI digested λ DNA marker; Lane 3, PCR validation of final knockout construct using P1 and P3 as primers. (F-H) PCR confirmation of pJAZZPbMPC2_c19e07 parental, intermediate, and final knockout constructs. F) PCR validation of the parental construct using P7 & P13 primer pair (lane 1, negative control) and P7 & P8 primer pair (lane 2). Lane 3, 1 kbp DNA ladder; lanes 4 and 5, PCR validation of the final construct using P7 & P13 primer pair and P7 & P8 primer pair (negative control), respectively. G)

Confirmation of the intermediate construct. Lane 1, 1 kb ladder; lane 2, PCR using P7 and P26 primer pair. H) XbaI restriction digestion of the intermediate construct (lane 1) and the final knockout construct (lane 3). Lane 2, 1 kbp ladder. Restriction digestion resulted in the release of DNA fragments of the expected sizes and confirmed the successful generation of constructs. kb in all panels should be read as kbp (kilo base pairs)

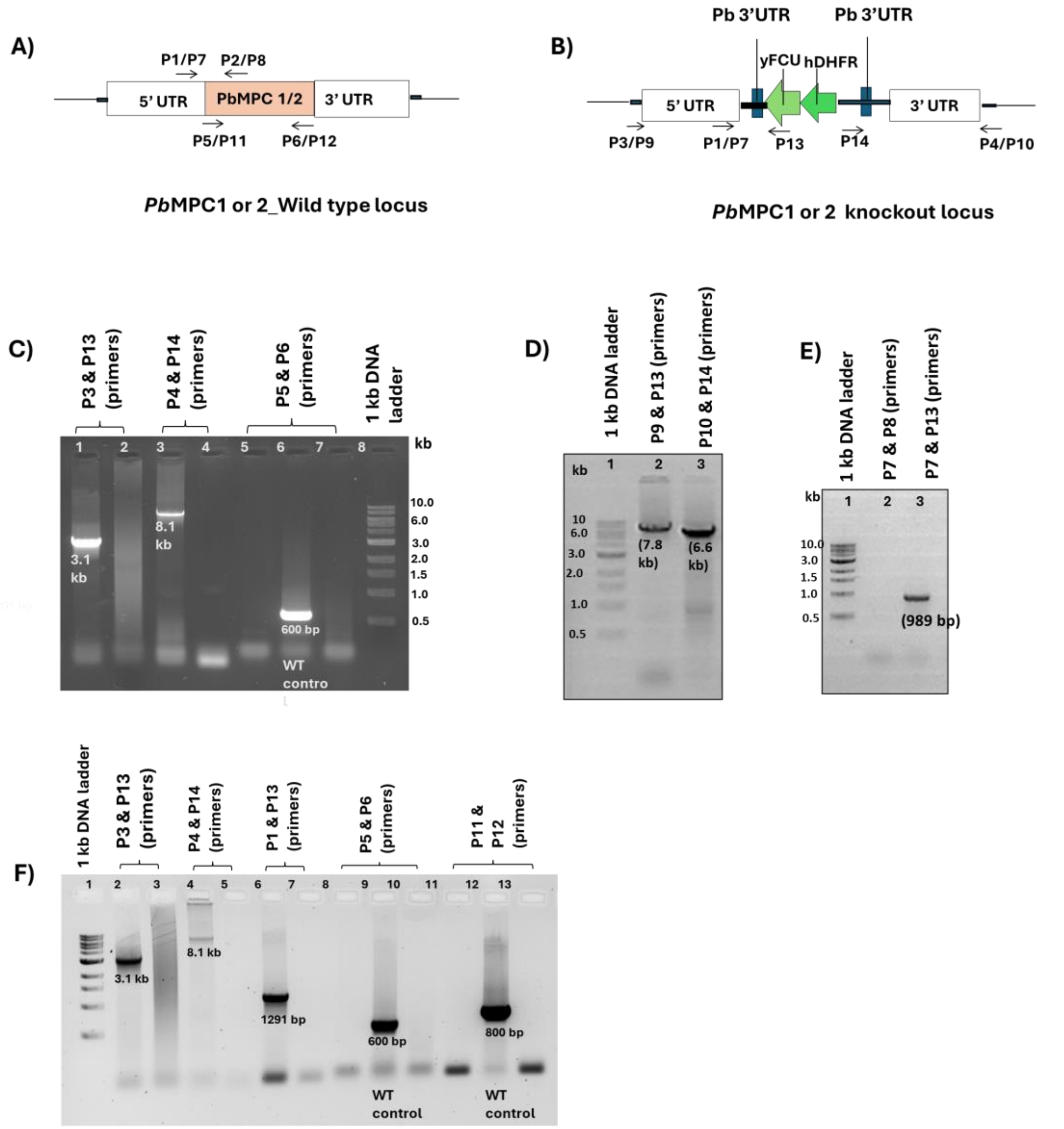

**Figure S6. Generation of Δmpc1, Δmpc2, and Δmpc1Δmpc2 parasites and confirmation by genotyping.** (A) and (B) Schematics of the MPC1 or 2 gene loci in the genome of *P. berghei* wild-type parasites and of the locus after knockout, respectively. Primers used for PCR genotyping are indicated. (C) Genotyping of *Δmpc1* parasites grown in C57BL/6 mice. PCR with primers P3 and P13, to confirm 5' integration using genomic DNA (gDNA) of *Δmpc1* parasites (lane 1) and without template (lane 2). PCR with primers P4 and P14, to confirm 3' integration using gDNA of *Δmpc1* parasites as template (lane 3) and without template (lane 4). PCR with P5 and P6 to confirm the absence of *mpc1* gene using gDNA of *Δmpc1* parasites (lane 5) as template. Genomic DNA from WT (lane 6), and without template served as positive and negative (lane 7) controls, respectively. 1kbp DNA ladder (lane 8). (D) Genotyping of drug-resistant *Δmpc2* parasites grown in C57BL/6 mice. Lane 1, 1kbp DNA ladder; lane 2, PCR with primers P9 and P13 to confirm 5' integration; lane 3, PCR with primers P10 and P14 to confirm 3' integration. (E) Lane 1, marker. PCRs with primers P7 and P8 (lane 2) confirm the absence of *mpc2* gene and with primers P7 and P13 (lane 3) confirm the presence of drug-resistant cassette using gDNA from *Δmpc2* parasites. The amplified products obtained by PCR are of the expected size. F) Genotyping of drug-resistant *Δmpc1Δmpc2* transfectants grown in C57BL/6 mice. Lane 1, 1 kb marker; lane 2, PCR with gDNA from transfectants for confirmation of 5' integration using P3 and P13 primers and lane 3, without template; lane 4, PCR confirmation for 3' integration using P4 and P14 as primers and lane 5, without template; lane 6 and lane 7, PCR confirmation for the presence of selection marker using P1 and P13 as primers and without template, respectively; PCR with primers P5 and P6 for the absence of *mpc1* gene using gDNA of *Δmpc1Δmpc2* parasites (lane 8), WT (lane 9) and without template (lane 10). PCR with primers P11 and P12 for the absence of *mpc2*

gene using gDNA of *Δmpc1Δmpc2* parasites (lane 11), WT (lane 12), and without template (lane 13). All the amplified DNA fragments obtained by PCR are of the expected size. The marker-free *Δmpc2* clone 1 was used further for genetic manipulation to generate *Δmpc1Δmpc2* parasites. kb in figure is kbp (kilo base pair)

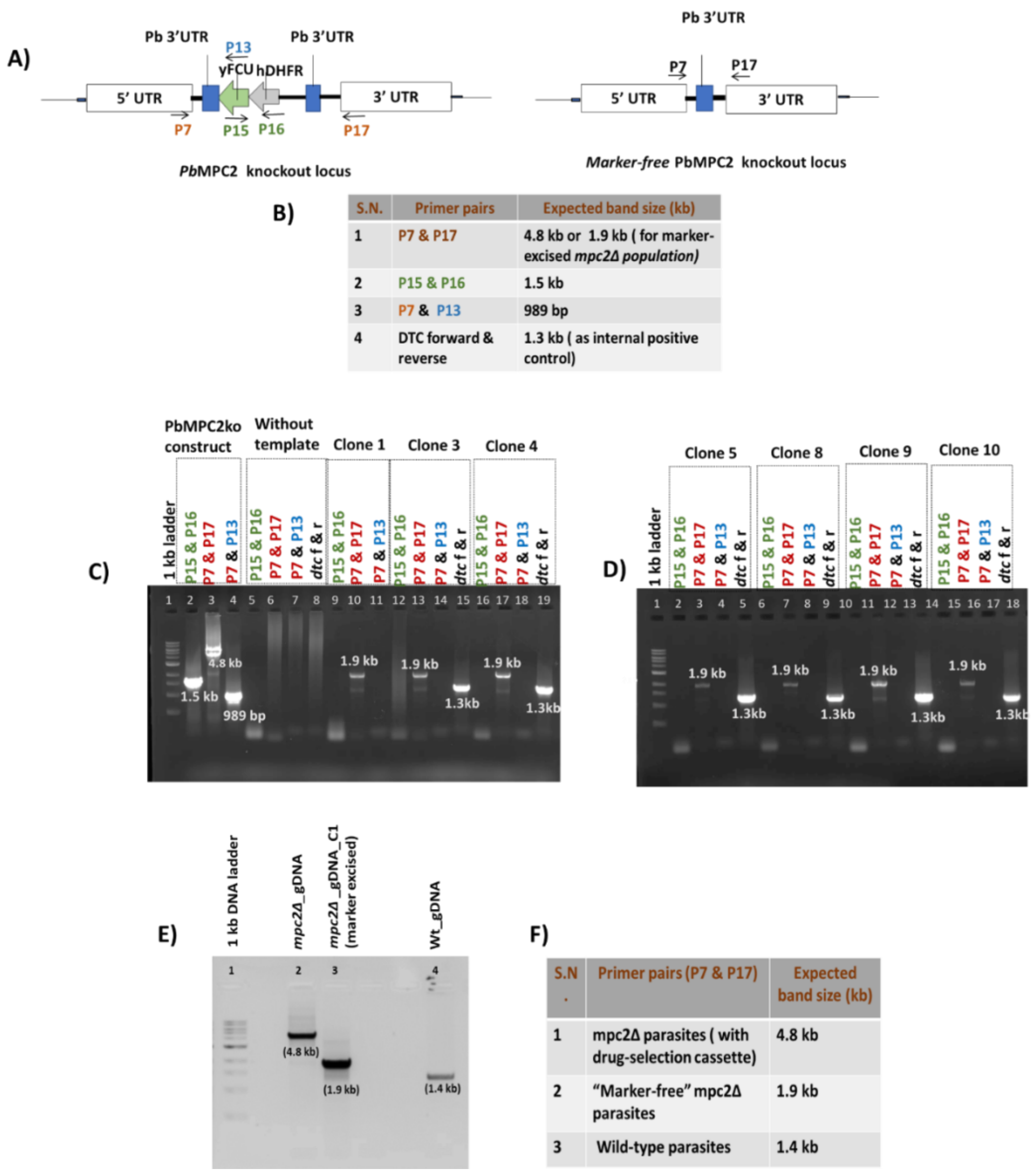

**Figure S7. Genotyping of marker-excised Δ*mpc2 P. berghei* knockout line.** A) Left panel, schematic of the locus in the genome of *Δmpc2 P. berghei* harbouring the hDHFR-yFCU cassette; right panel, schematic of the locus in the genome of marker excised *Δmpc2 P. berghei* lacking the hDHFR-yFCU cassette. The positions of different primers are indicated as arrows. B) Table listing the expected amplified DNA fragment size for specific pairs of oligonucleotide primers. (C-D) PCR genotyping for the confirmation of marker excision in *mpc2* knockout *P. berghei* clones obtained by limited dilution cloning. The seven positive clones were obtained (C1, C3, C4, C5, C8, C9 and C10). The marker-free *Δmpc2* clone 1 was used for further study. C) Lane 1, 1 kbp ladder; lanes 2,3 and 4, PCR with *Δmpc2* final knockout construct DNA as a template (positive control) with three different sets of oligonucleotides; yFCU (P15) forward and hdhfr (P16) reverse, P7 forward and P17 reverse, P7 forward and P13 reverse primers, respectively. Lanes 5, 6, 7, and 8, PCR without template (negative control) with four different sets of oligonucleotides; P15 forward and P16 reverse, P7 forward and P17 reverse, P7 forward and P13 reverse and Pbdtc forward and reverse primers, respectively. Lane 9- lane 11; lane 12- lane 15; lane 16- lane 19 show PCR results of clone 1, clone 3, and clone 4 using P15 forward and P16 reverse, P7 forward and P17 reverse, and P7 forward and P13 reverse primers, respectively. D) Lane 1, 1 kbp ladder; lane 2- lane 5; lane 6- lane 9; lane 10-lane13; lane 14-lane 18, show PCR results of clone 5, clone 8, clone 9, and clone 10 using P15 forward and P16 reverse, P7 forward and P17 reverse, and P7 forward and P13 reverse primers, respectively. Pbdtc forward and reverse primers serve as internal positive control. (E-F) Revalidation by PCR of marker-free *Δmpc2 P. berghei* clone 1. Lane 1, 1 kbp ladder; lane 2, PCR with gDNA from *Δmpc2 P. berghei* as template; lane 3, PCR with gDNA from marker-excised *Δmpc2* clone 1 as template; lane 4, PCR with gDNA from wild-type *P. berghei* as template. Primer pair: P7 forward and P17 reverse were used for PCRs. The PCR-amplified bands of the expected size were observed. kb is kbp (kilo base pair).

A)

| Expected Size (Transcript length) | Expected Size (Genome sequence length) |
|---|---|
| PbMPC1 (368 bp) | PbMPC1 (547 bp) |
| PbMPC2 (396 bp) | PbMPC2 (753 bp) |
| PbGAPDH ( 1 kb) | PbMPC2 ( 1.2 kb) |

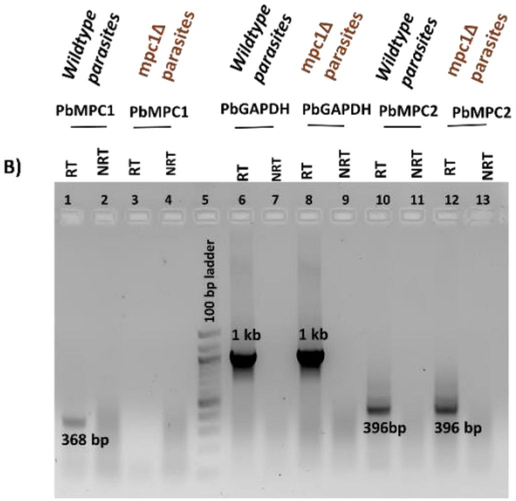

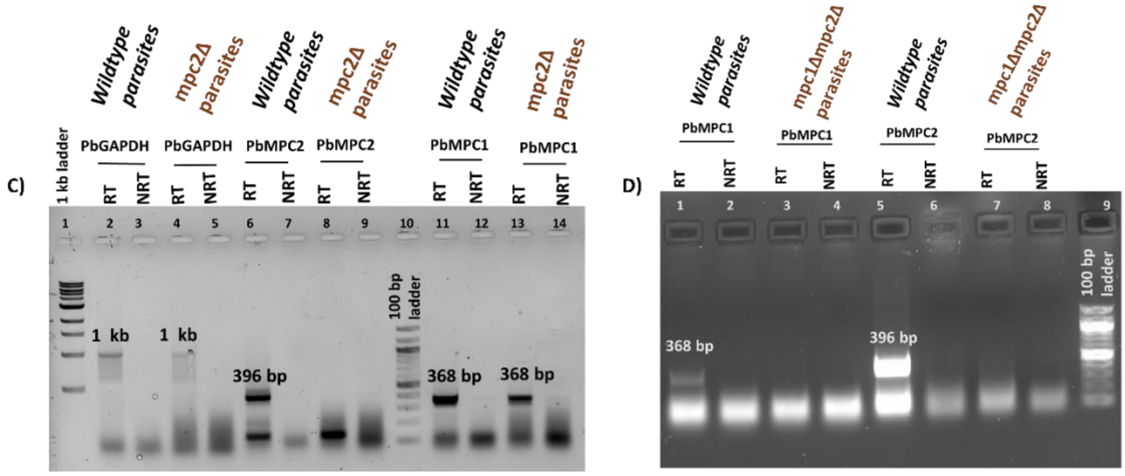

**Figure S8. RT-PCR confirmation for the absence of transcripts for MPC1 or MPC2 or both in *Δmpc1*, *Δmpc2*, and *Δmpc1Δmpc2* parasites.** A) Table lists the sizes of DNA fragments expected from PCR using either gDNA or cDNA as template and *P. berghei mpc1*, *mpc2*, or *gapdh* forward and reverse primers. B) Confirmation for the absence of *mpc1* gene expression at the transcript level by RT-PCR. Lanes 1 and 3, RT-PCR using total RNA from *mpc1* knockout and wild-type parasites, respectively using gene-specific *mpc1* forward (P5) and reverse (P6) primers. PCR with no reverse transcriptase (NRT) served as negative control (lanes 2 and 4). Lane 5, 100 bp marker; lane 6 - lane 13, RT-PCR using total RNA from *mpc1* knockout and wild-type parasites with gene-specific *gapdh* (P53 and P54) and *mpc2* (P11 and P12) primers, as internal positive controls. C) Confirmation for the absence of *mpc2* gene expression at the transcript level by RT-PCR. Lanes 1 and 10, 100 bp ladder. Lanes 6 and 8, RT-PCR with RNA isolated from *mpc2* knockout parasites using gene-specific *mpc2* forward (P11) and reverse (P12) primers. PCR with no reverse transcriptase (NRT) served as negative control (lanes 7 and 9). Lane 2-5 and lane 11-14, RT-PCR with *mpc2* knockout and wild-type parasites using gene-specific *gapdh* (P53 and P54) and *mpc1* (P5 and P6) primers, as internal positive controls. D) RT-PCR results from *Δmpc1Δmpc2* parasites. RNA was isolated from wild-type (positive control) and *Δmpc1Δmpc2* parasites. RT-PCR was carried out with *mpc1* gene-specific primer pair, P5 and P6 (lane 1-4), and *mpc2* gene-specific primer pair, P11 and P12 (Lanes 5-8) using wild-type or *Δmpc1Δmpc2* cDNA as template. Lane 9, 100 bp ladder. RT-PCR results confirmed the expression of *mpc1* and *mpc2* genes in wild-type but not in *Δmpc1Δmpc2* parasites. Note, RT-PCR experiments were done on the clonal line of *Δmpc2* and *Δmpc1Δmpc2* while in the case of *Δmpc1* an uncloned population was used.

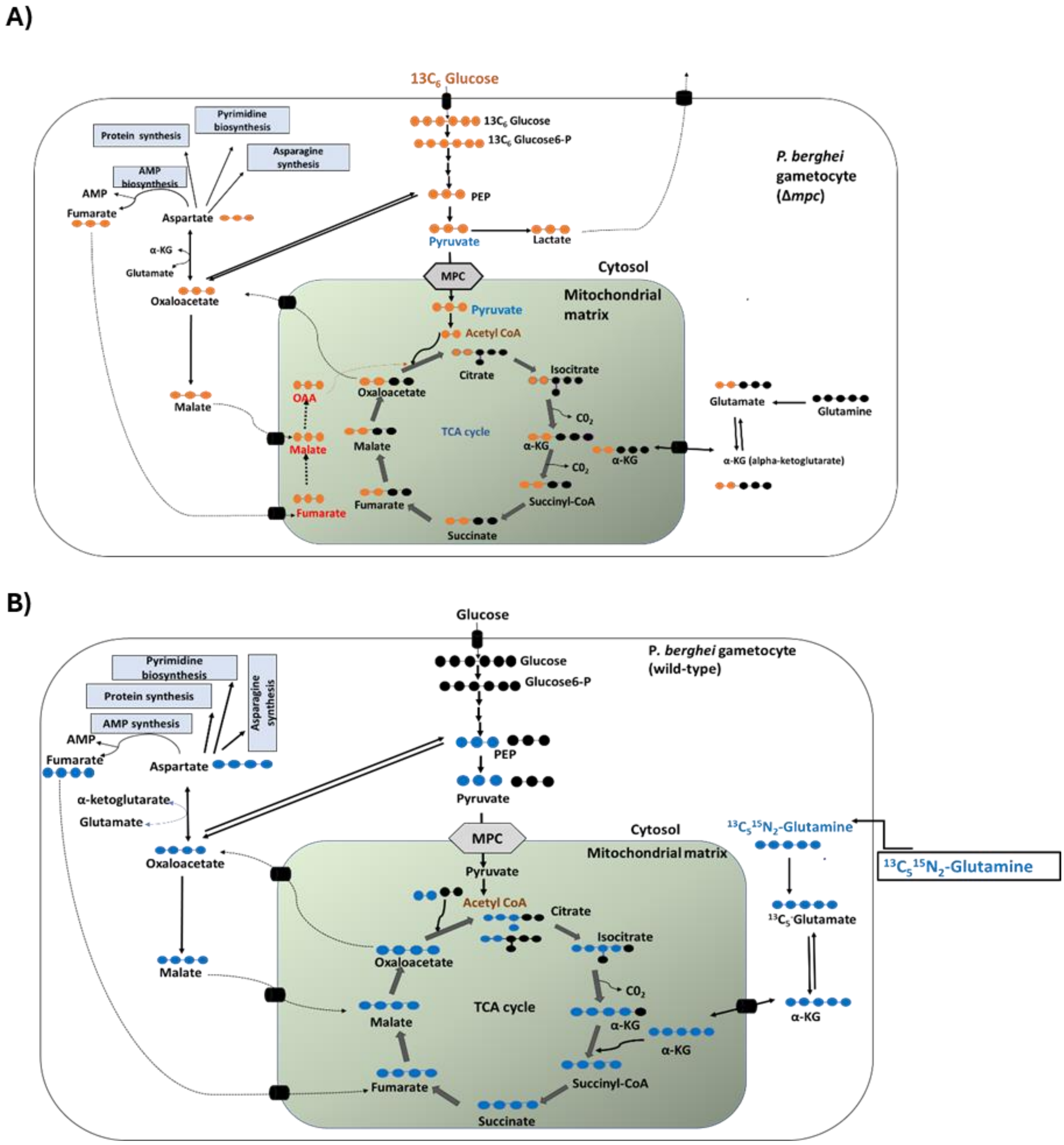

**Figure S9. The metabolic flux of $^{13}$C-carbons through glycolysis and TCA cycle in the presence of U-$^{13}C_6$ glucose as carbon source and U-$^{13}C_5{}^{15}N_2$ glutamine as carbon and nitrogen source. A)** In the cytosol, all glycolytic intermediates should display either M+6 or M+3 labelling. This figure highlights the flux of $^{13}$C-carbons through the first turn of the TCA cycle. The black-filled circles indicate $^{12}$C-atoms present in a metabolite while orange-filled circles indicate $^{13}$C-atoms. Anaplerotic reactions that feed the TCA cycle should display M+3 labelling of intermediates. $^{13}C_6$-glucose-derived labelled glutamate may be seen at low levels. B) In the cytosol, glutamine is first converted to glutamate which is then further metabolized to alpha-ketoglutarate to fuel the TCA cycle within the mitochondrion. The intermediates generated through the TCA cycle should show M+4 labelling. Very low levels of labelled glycolytic intermediates such as PEP and pyruvate could enter the TCA cycle and show M+2 labelling of TCA cycle intermediates. The black-filled circles indicate $^{12}$C-atoms present in a metabolite while blue-filled circles indicate $^{13}$C-atoms.

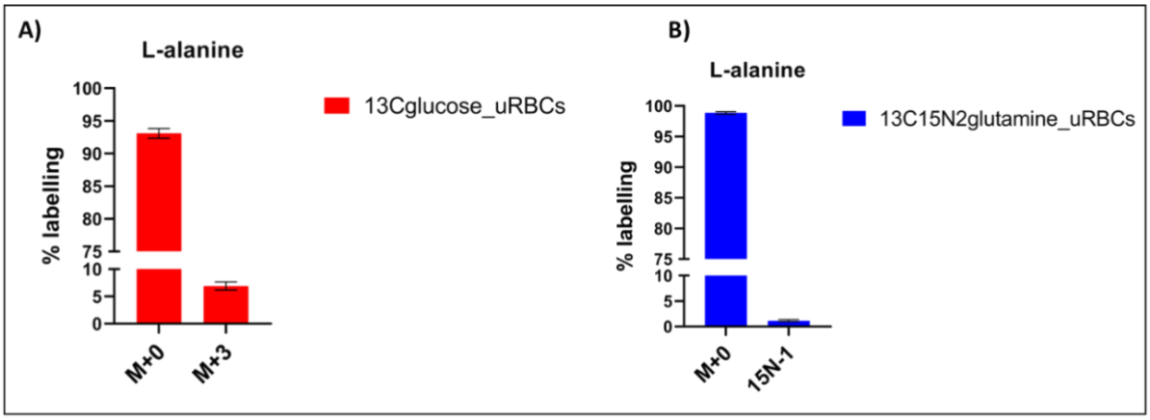

**Figure S10. $^{13}$C- and $^{15}$N- enrichment of L-alanine extracted from uninfected erythrocytes (uRBCs).** Erythrocytes were incubated for 90 minutes with $^{13}C_6$-glucose (A) and $^{13}C_5{}^{15}N_2$-glutamine (B). $^{13}$C or $^{15}$N enrichment of alanine extracted from erythrocytes is plotted, with the y-axis representing the percent abundance of isotopologues of alanine resulting from $^{13}C_6$-glucose and $^{13}C_5{}^{15}N_2$-glutamine labelling. The graphs for percentage labelling of alanine were plotted after correcting for natural abundance (NA) using IsocorrectoR. Error bar shows SD of N=3 technical replicates from one experiment.
